# Prioritizing Maize Transcriptional Regulators through Multi-Omic Network Integration

**DOI:** 10.1101/2024.02.26.582075

**Authors:** Fabio Gómez Cano, Jonas Rodriguez, Peng Zhou, Yi-Hsuan Chu, Erika L. Ellison, Lina Gomez-Cano, Arjun Krishnan, Nathan M. Springer, Natalia de Leon, Erich Grotewold

## Abstract

Transcription factors control all biological processes, yet most remain functionally uncharacterized. As in many crops, maize functional genomics datasets are typically generated independently across genotypes, tissues, and laboratories, with little sample overlap and uneven transcription factor coverage, limiting methods that require matched measurements to make functional inferences. Here, we asked which gene-level integration strategy best predicts transcription factor function from such data. We assembled four transcription factor–target gene network layers from 46 expression-based regulatory networks, 283 protein– DNA interaction assays, and expression quantitative trait loci mapped with 16 million single-nucleotide polymorphisms across 304 inbred lines, comprising ∼4.6 million interactions, and benchmarked three integration strategies against published knockout transcriptomes. Predicted Gene Ontology terms were more similar to knockout responses than expected by chance, and summarizing each gene’s connectivity pattern across the combined network as a numerical representation (a gene embedding) achieved the highest similarity where predictions existed, annotated more than eightfold more transcription factors (2,910), remained informative when data layers were missing, and produced significant terms in only ∼12% of randomized networks, versus ∼28% and ∼72% for two overlap-based strategies. The resulting catalog recovers known regulators, prioritizes candidate regulators of hormonal, developmental, and metabolic processes across conditions, and identifies candidate redundant or diverged paralogs, providing a first testable function for thousands of transcription factors that lack binding or mutant data. Because it uses only data types now available for many crops, the strategy transfers directly to other plant species.

**Article Summary:** Transcription factors are proteins that control gene expression, shaping crop traits from development to the production of valuable compounds. Yet most maize transcription factors have no known function and no characterized mutants. Gomez-Cano et al. integrated gene expression, protein–DNA binding, and genetic variation into one network to predict the functions of about 2,900 transcription factors, most of which had no binding or mutant data. In published mutants, predicted functions matched observed changes in gene activity better than expected by chance. The catalog, networks, and code are publicly available and prioritize regulators of hormonal, developmental, and metabolic processes for experimental testing.

## Introduction

Like other organisms, plants use highly interconnected molecular networks that coordinate every cellular process, from cell division to metabolite synthesis. Among these, gene regulatory networks play central roles by controlling the expression of essentially every gene, a function carried out mainly by transcription factor (TF) proteins (Lambert *et al*. 2018). Transcriptional regulation requires direct protein–DNA interactions (PDIs) between TFs and specific *cis*-regulatory elements located in proximal (promoters) or distal (enhancers/silencers) regions of their target genes (Schmitz *et al*. 2022). Collectively, sets of PDIs form interconnected gene regulatory networks (GRNs). The architecture of GRNs is commonly survey with gene-centered approaches, such as yeast one-hybrid assays (Arda and Walhout 2010; Mejia-Guerra *et al*. 2012; Yang *et al*. 2016), and TF-centered approaches, such as chromatin immunoprecipitation sequencing (ChIP-seq) for *in vivo* PDIs and DNA affinity purification sequencing (DAP-seq) for *in vitro* PDIs (Yang *et al*. 2016; O’Malley *et al*. 2016). These categories are useful but not absolute. ChIP-seq scales poorly across many TFs, whereas DAP-seq captures interactions outside the native chromatin context (Yang *et al*. 2016). GRN organization has implications for phenotypic variation (Deplancke *et al*. 2006), responses to abiotic and biotic stress (Nakashima *et al*. 2014; Birkenbihl *et al*. 2017; Sun *et al*. 2022), development (Marand *et al*. 2023), speciation (Mack and Nachman 2017), and adaptation and diversification (Mack and Nachman 2017; Bowles *et al*. 2020; Marand *et al*. 2023). Consequently, understanding GRN structure and dynamics therefore offers a valuable route to manipulating desired phenotypes.

As in other plant and animal systems, agronomically important traits in maize are quantitative, arising from multiple loci which independently explain small fraction of trait variance (Riedelsheimer *et al*. 2012; Wen *et al*. 2014, 2016; Xiao *et al*. 2017; Mazaheri *et al*. 2019; Zhou *et al*. 2019). This genetic architecture underlies a valuable metabolic diversity (Riedelsheimer *et al*. 2012; Wen *et al*. 2014, 2016; Zhou *et al*. 2019) built on a large genetic diversity (Schnable *et al*. 2009; McMullen *et al*. 2009; Hufford *et al*. 2021) modulated by organ identity, developmental stage, and environment cues (Wen *et al*. 2014; Kusmec *et al*. 2017; Xiao *et al*. 2017). However, because many genes and regulatory elements jointly shape each trait, locus–trait associations alone rarely resolve the underlying mechanisms.

A distinctive feature of the maize genome is a recent whole-genome duplication ∼5–12 Mya, making maize an ancient allotetraploid (Wei *et al*. 2007). Maize is rich in tandem-duplicated genes (Kono *et al*. 2018) and highly enriched in transposable elements (∼85% of the genome) (Schnable *et al*. 2009). This duplication produced two subgenomes (maize1 and maize2) with unequal gene loss and expression (Schnable *et al*. 2011). Notably, the dominant subgenome, maize1, contributes more to phenotypic variation (Renny-Byfield *et al*. 2017). However, the mechanisms driving this asymmetry remain unclear, though co-expression and multi-network studies point to divergence across transcriptional, signaling, and interaction networks (Li *et al*. 2016; Han *et al*. 2023), with direct implications for trait prioritization and maize evolution.

The rapid expansion of multi-omic datasets has enabled characterization of the genetic and phenotypic complexity of biological systems (Tolani *et al*. 2021). Approaches for integrating these datasets continue to evolve and can be broadly classified as conceptual integration based on overlapping observations, statistical integration, network-based integration, and machine-learning-based integration (Depuydt *et al*. 2023). Many widely used methods, including multi-omics factor analysis (Argelaguet *et al*. 2018) and similarity network fusion (Wang *et al*. 2014), assume that different data types are measured in the same samples, as is common in patient cohorts. Gene-level integration provides an alternative by linking datasets through shared genes and therefore does not require matched samples. Network-based approaches, in particular, can recover gene function from heterogeneous datasets by merging distinct molecular networks into a single map and summarizing each gene’s overall pattern of connectivity as a numerical representation (an embedding), placing genes that share regulatory partners close together even when individual datasets are incomplete (Cho *et al*. 2016).

In maize, technological advances have generated extensive functional genomics resources, including TF-binding profiles (Galli *et al*. 2018; Ricci *et al*. 2019; Tu *et al*. 2020), accessible chromatin regions (Rodgers-Melnick *et al*. 2016; Ricci *et al*. 2019; Marand *et al*. 2021), expression and co-expression atlases (Sekhon *et al*. 2011; Stelpflug *et al*. 2016; Hoopes *et al*. 2019; Zhou *et al*. 2020), and population-level transcriptomic, proteomic, and metabolic data (Li *et al*. 2013; Wen *et al*. 2014, 2016; Kremling *et al*. 2018; Mazaheri *et al*. 2019; Zhou *et al*. 2019; Shrestha *et al*. 2022). These resources have motivated efforts to integrate complementary molecular data (Wen *et al*. 2016, 2018; Liu *et al*. 2016; Walley *et al*. 2016; Jin *et al*. 2017; de Abreu e Lima *et al*. 2018; Schaefer *et al*. 2018; Lee *et al*. 2019; Yang *et al*. 2022), yet they present a challenge common to crops, as most datasets were generated independently across genotypes, tissues, and laboratories with little sample overlap and uneven TF coverage. For example, the PDI datasets reanalyzed here cover only 142 TFs, and published loss-of-function transcriptomes span a reduce list TFs, leaving most maize TFs without experimental data to guide prioritization. How best to integrate these unmatched, incomplete gene-level data layers has not been systematically evaluated.

In this study, we analyzed genetic and gene-expression variation across 304 maize inbred lines. We reanalyzed over 300 publicly available ChIP- and DAP-seq experiments together with 45 previously analyzed random-forest regulatory networks (RFNs, expression networks based on decision-tree models that identify which TFs best predict each target gene’s expression across samples) (Zhou *et al*. 2020) to evaluate how accurately different integration methods recover TF’s function annotations. Specifically, we constructed four molecular network layers from these data and compared three integration strategies for predicting TF function from their inferred target genes. Merging all four layers into a single weighted network and summarizing each gene’s connectivity pattern as a numerical representation (a gene embedding) produced the strongest overall predictions while retaining information from layers with incomplete TF coverage. We benchmarked these predictions against published knockout transcriptomes for 14 TFs, randomized networks, and experimentally determined yeast one-hybrid interactions. The resulting catalog recovered known hormonal, developmental, and metabolic regulators, prioritized new candidates, resolved condition-specific TF activity across tissues and genotypes, and combined embeddings with sequence and expression data to identify candidate redundant or diverged TF paralogs. We release the catalog, network layers, and gene embeddings (Data Availability) as a resource for prioritizing maize transcriptional regulators.

## Results

### Construction of a multi-layer maize regulatory network

To predict TF functions across diverse biological contexts, we assembled four complementary network types, associating TFs to candidate target genes, using experimental evidence from three categories of publicly available data (**Fig 1A**). Specifically, we used expression data to generate random-forest regression regulatory networks (RFNs). Genotype and expression data for 304 maize inbred lines (**Supplementary Table 1**) were used to define the *trans*-eQTL gene-association network (hereafter, the *trans*-eQTL network). Protein–DNA interaction data from DAP-seq and ChIP-seq (**Supplementary Table 2**) define the protein–DNA interaction (PDI)-based gene regulatory network (GRN) and, combined with *cis*-eQTLs, a *cis*-eQTL-supported GRN. Each layer is described below and detailed in Methods, and their sizes are summarized in **Fig 1B**.

**Fig 1.**
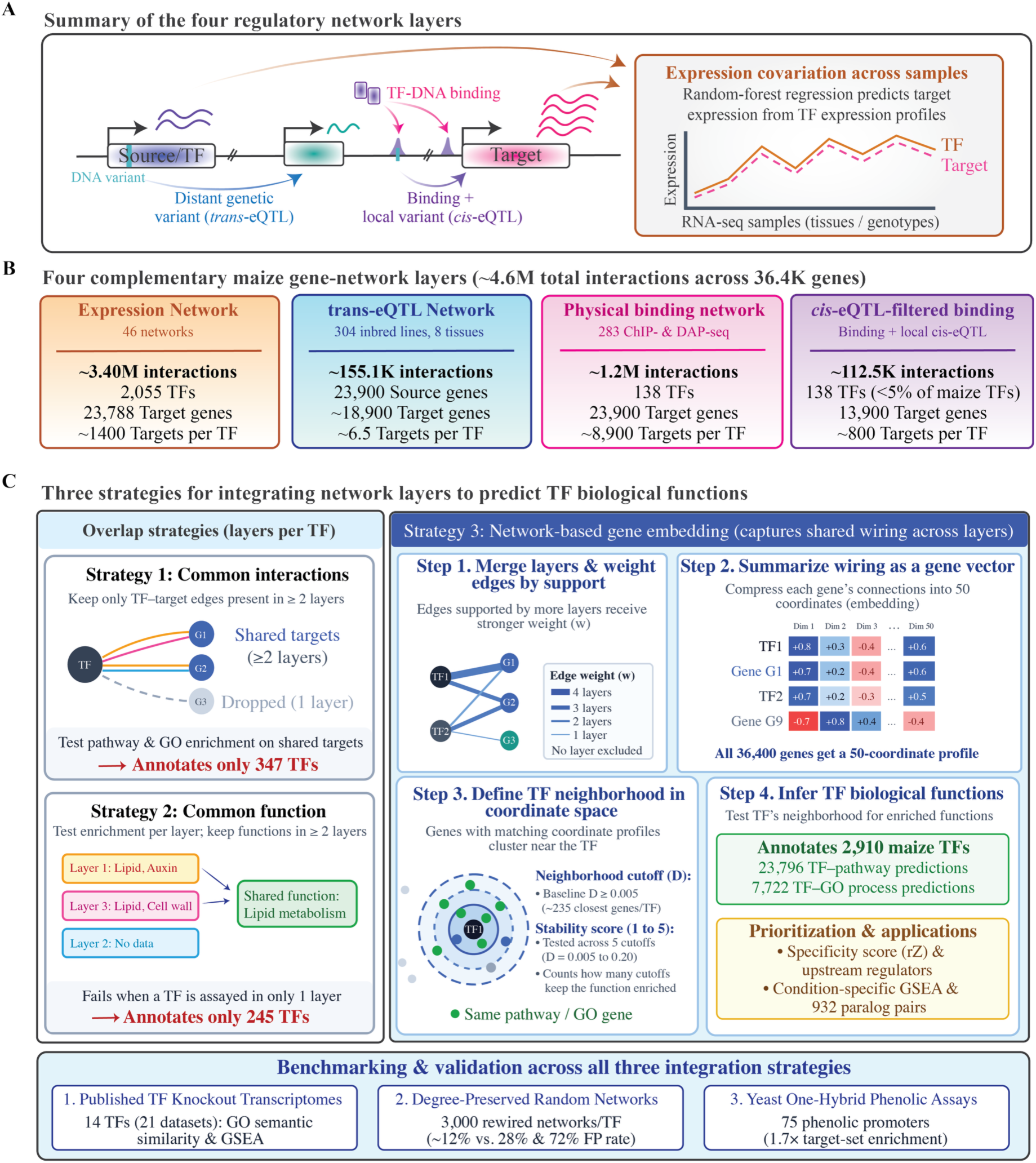
Construction and integration of a multi-layer maize gene regulatory network to predict transcription factor (TF) functions. (A) Four types of biological evidence linking regulatory source genes and TFs to candidate target genes: (1) expression covariation across RNA-seq samples modeled by random-forest regression, (2) distant genetic variation (*trans*-eQTLs) associated with target gene expression across 304 inbred lines, (3) physical TF–DNA binding measured by ChIP-seq and DAP-seq at promoter (≤3 kb) or distal sites overlapping accessible chromatin, and (4) physical TF binding sites supported by a local variation *cis*-eQTL. (B) Summary metrics of the four layers. (C) Comparison of three integration strategies for predicting TF biological roles. We evaluate all three strategies against published TF knockout transcriptomes, degree-preserved randomized networks, and yeast one-hybrid assays.

### Random-forest regulatory networks (RFNs)

We started from 45 previously published expression-based regulatory networks (Zhou *et al*. 2020) and added one network built from expression data for the 304 Wisconsin Diversity panel inbred lines (Mazaheri *et al*. 2019), following the same procedure used before (Zhou *et al*. 2020) (*Methods*, Supplementary Table 3). Briefly, for each target gene, random-forest regression models its expression as a function of TF expression across samples, generating more robust TF–target associations than classic pairwise correlations because it evaluates all candidate TFs simultaneously and captures nonlinear and combinatorial effects. Each network was restricted to maize genes syntenic to *Sorghum bicolor* genes (Zhang *et al*. 2017) to limit the effect of pseudogenes, which inherit GO and pathway annotations from their functional paralogs and would otherwise bias the enrichment results. The same filter was applied in all enrichment analyses below. On average, each RFN contained 1,055 TFs (**Supplementary Figure 1A**) with ∼74 predicted targets each (**Supplementary Figure 1B**). The union of all 46 RFNs yielded ∼3.4 million associations involving 2,055 TFs and 23,788 target genes (∼1,440 targets per TF in the merged network; **Supplementary Figure 1C**).

### *Trans*-eQTL network

To build the *trans*-eQTL network, we identified eQTLs across eight tissues spanning several developmental stages (*Methods*). Across tissues, we tested 15.5–16.7M SNPs against the expression of 15.3K–26.4K genes (**Supplementary Table 4**) and retained an initial set of ∼6.7M eQTL– gene associations after removing non-significant eQTLs, non-syntenic genes, and trans associations falling outside ±100 kb of the target gene (**Supplementary Figure 2A**). Genes colocalized with eQTLs were defined as "source genes," and their corresponding associated transcripts as "target genes." After collapsing redundant associations, the *trans*-eQTL network held ∼155K associations across 23.9K source and 18.9K target genes. We sort source and target genes in six functional classes (TFs, co-regulators, Mediator subunits, kinases, enzymes, and other; Supplementary Table 5) to describe its representation on the network. After normalizing by *trans*-eQTL SNPs per class, interaction rates were uniform across classes (0.11–0.15 interactions per SNP, with 96–100% of SNP-bearing genes acting as sources), indicating that class contributions reflect SNP frequency rather than excess connectivity (**Supplementary Figure 2B–2D**). However, despite the lack of enrichment for a specific class, the *trans*-eQTL network recovered interactions consistent with both TF–target regulation and protein–protein interactions. For example, HSF20 showed 354 targets, including 27 previously reported heat-response genes (Zhou *et al*. 2022) and five known HSF20 physical interactors (**Supplementary Table 6**) (Zhu *et al*. 2016).

### PDI-based GRN and *cis*-eQTL-supported GRN

We reanalyzed 283 TF-centered PDI experiments for 142 TFs (captured in 245 ChIP-seq and 38 DAP-seq) through a single pipeline to control for technical variation (*Methods*). We observed, on average, 52K binding peaks per TF (∼7.6M PDIs total, **Supplementary Table 7**, **Supplementary Figure 3A**). We retained high-confidence peaks by (i) keeping only peaks overlapping accessible chromatin regions from a single-nucleus ATAC-seq atlas (Marand *et al*. 2021) and (ii) removing low-coverage peaks (normalized read coverage Z-score ≤ −0.5) (*Methods*), leaving ∼3.4M high-confidence peaks (**Supplementary Figure 3B–3D**). Retained peaks were assigned to a target gene by their distance to the nearest annotated TSS, treating peaks within 3 kb as proximal and peaks between 3 kb and 50 kb as distal (**Supplementary Figure 3E**, *Methods*). Proximal peaks (≤ 3 kb from TSS) were assigned to the nearest gene directly, whereas distal peaks were kept only when independently supported by a *cis*-eQTL. Thus, GRN comprises proximal PDI-mapped promoters, while the *cis*-eQTL-supported GRN is defined by distal binding plus local cis-eQTL support. Overall, the two layers contained ∼1.2M (GRN) and ∼112.5K (*cis*-eQTL-supported GRN) TF–target interactions involving 138 TFs and ∼23.9K and ∼13.9K target genes, respectively.

### TF functional annotation

A key difference among the four layers is how many TFs each contains and how many target genes those TFs have. Only 111 TFs are present in all four layers with at least one target gene, and only 17 TFs have at least ten targets in every layer (**Supplementary Figure 4A–C**). This is partly due to the *trans*-eQTL network, in which each source gene has few targets (on average, ∼6.5). To evaluate the effect of this sparsity on TF’s functional annotation, we implemented and evaluated three strategies for assigning functional roles to each of the TFs on the corresponding layers, which we call *common interactions*, *common function*, and *network-based*. In each case, we tested for enrichment of metabolic pathways (PWYs) and Gene Ontology (GO) terms among the targets of each TF (**Fig 1C**, *Methods*). *Common interactions*, the most conservative, assumes that only TF–target interactions shared across layers reflect true targets. *Common function* prioritizes functions enriched consistently across layers. Finally, the network-based strategy merges all four layers into a single weighted network and summarizes each gene’s overall connectivity pattern as a 50-value numerical profile (a gene embedding; **Fig 1C**). Because two genes receive similar embeddings whenever they share direct or indirect regulatory partners across layers, we can define a functional neighborhood of the most closely connected genes around every TF and test that neighborhood for PWY and GO enrichment (*Methods*). Unlike the other two strategies, this approach can annotate any TF present in the combined network even when individual layers are sparse or lack binding data for that TF.

Extended descriptions of the *common interactions* and *common function* strategies are provided in **Supplementary File 1**. Briefly, *common interactions* strategy identified 11,362 significant TF–process associations (2,812 TF–PWY and 8,550 TF–GO, **Fig 2A**), averaging ∼8 PWYs and ∼70 GO terms per annotated TF (**Fig 2B**), and annotated 347 TFs in total (**Supplementary Table 8**, **Fig 2C**). The *common function* strategy yielded 7,081 associations (727 TF–PWY and 6,354 TF–GO, **Fig 2A**), averaging ∼3.6 PWYs and ∼58 GO terms per annotated TF (**Fig 2B**), and annotated 245 TFs (**Fig 2C**). In comparison, the network-based strategy annotated over eight times more TFs (**Fig 2A–C**) and is detailed in the following section.

**Fig 2.**
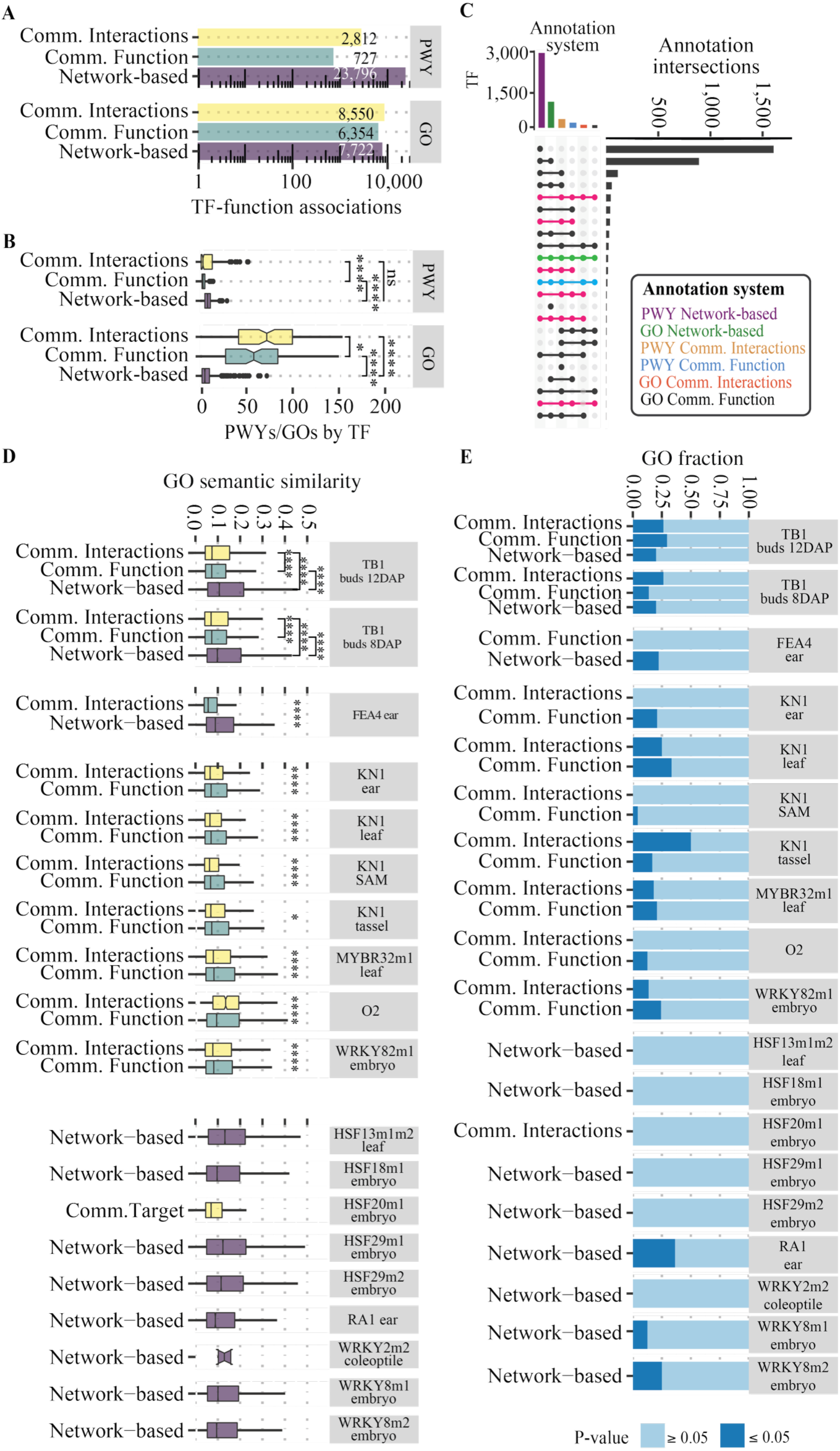
Benchmarking of the three integration strategies against knockouts and random networks. (A) Total PWYs and GO terms predicted per strategy after combining all TFs. (B) PWYs and GO terms per TF. (C) UpSet plot of TFs annotated per strategy and annotation system (fuchsia, PWY; blue, GO; green, both; black, annotated by ≥ 1 method). (D) GO semantic similarity between predicted and knockout GO terms, for the top-10 most-similar terms per TF and strategy. (E) Fraction of predicted GO terms significantly enriched (GSEA) in the corresponding knockouts. Asterisks: *P ≤ 0.05, **P ≤ 0.01, ***P ≤ 0.001, ****P ≤ 0.0001 (two-sided t-test). “TFm” denotes multiple mutant lines for the same TF.

### Network-based integration annotates the most TFs

To combine all four layers into a single weighted network, we weight each interaction by the number of layers supporting it, and developed a low-dimensional embedding for every gene (**Supplementary Figure 6A**, *Methods*). The combined network comprised 4.6M interactions across 36.4K genes. Unlike the other two strategies, this assigned a 50-value embedding to every gene in the combined network, including TFs present only in the RFN and *trans*-eQTL layers, allowing TFs without direct binding (PDI) data to still be evaluated. To define each TF’s functional neighborhood (the set of genes most closely connected to that TF in the integrated network), we ranked all genes by how similar their embeddings were to the TF’s embedding and retained those passing a neighborhood similarity cutoff (D ≥ 0.005, where D scales from 0 for unrelated genes to 1 for identical profiles; *Methods*). At this default cutoff, each TF’s functional neighborhood contained an average of 235 genes, which we then tested for PWY and GO enrichment to assign functional predictions to the TF (**Supplementary Figure 6A, B**).

We identified 23,796 TF–PWY and 7,722 TF–GO significant associations (FDR ≤ 0.1; **Fig 2A**, **Supplementary Table 8**), averaging ∼8 PWYs and ∼7 GO terms per TF (**Fig 2B**). In total, we found 2,910 TFs enriched in at least one function, and 1,030 TFs enriched for both PWYs and GO terms (**Supplementary Figure 6C**). These TFs span 82 families and account for ∼34% of the proteins in those families (**Supplementary Figure 6D**). Compared with the other two strategies, the *network-based* approach recovered far more TF–PWY associations (**Fig 2A**) but assigned the fewest GO terms per TF (**Fig. 2B**), while annotating over eight times as many TFs overall (**Fig 2C**). Of the 2,910 annotated TFs, fewer than 5% have any published binding data (the GRN layer contains 138 TFs), and about 1% have a published loss-of-function transcriptome (29 TFs, **Supplementary Table 9**). For the remainder, the catalog provides the first testable functional hypothesis.

Because the size of each TF’s functional neighborhood depends on the similarity cutoff D used to include neighbor genes, we evaluated how sensitive our predictions are to neighborhood size by repeating the enrichment analysis across five increasingly strict cutoffs spanning a 40-fold range (from the default D ≥ 0.005 down to D ≥ 0.20, which reduces the median neighborhood from 227 to 54 genes; *Methods*, **Supplementary File 3**). All 2,915 TFs retained annotatable neighborhoods at every threshold, and the number of significantly annotated TFs was stable even though median neighborhood size shrank four-fold (**Supplementary Figure 16A, 16B**), indicating that annotation yield is not an artifact of neighborhood size. Associations near the significance boundary turned over between thresholds, whereas the strongest associations were retained two to three times more often (**Supplementary Figure 16C**). To distinguish predictions that hold across neighborhood sizes from those that depend on a single cutoff, we report a threshold-stability score (ranging from 1 to 5, counting how many of the five D cutoffs support a given TF– process association) alongside the catalog (**Supplementary Figure 16D**). Specifically, 50.2% of TF–PWY associations were significant in at least three of five thresholds (**Supplementary Table 8**). Repeating this five-cutoff sweep for the TF–GO predictions yielded a similarly high consistency, with up to 52.0% of scorable TF–GO associations remaining significant across at least three of the five cutoffs, closely matching the stability observed for the metabolic pathway predictions (**Supplementary File 3**).

### Benchmarking against TF knockouts favors embedding-based integration

We next asked which integration strategy provides functional predictions that hold up against experimental evidence. We benchmarked all three strategies against published knockout RNA-seq data for 14 TFs (comprising 21 mutant × tissue datasets, **Supplementary Table 9**) (Zhou *et al*. 2020; Ellison *et al*. 2023), of which 13 yielded predictions that could be compared. We used two independent tests. First, the overlap between predicted PWYs/GO terms and those enriched among differentially expressed genes in each knockout. Second, gene-set enrichment analysis (GSEA) of the predicted terms within each knockout, ranked by log2 fold-change (Subramanian *et al*. 2005) (*Methods*).

Predicted metabolic pathways overlapped poorly with enriched metabolic pathways from knockout for every method (**Supplementary Figure 7**), whereas predicted and observed GO terms were more similar than expected by chance when evaluated by GO semantic similarity, which measures how closely related two GO terms are in the ontology hierarchy (P < 0.05, **Supplementary Figure 8**). The three strategies were not equivalent, with GO terms predicted by the *network-based* and *common function* strategies being significantly closer to the knockout GO terms than those from *common interactions* (P < 0.05, Wilcoxon test, **Fig 2D**). Notably, when predictions were available across methods, the *network-based* strategy achieved the highest semantic similarity (**Fig 2D**; TB1 and FEA4). However, the methods showed complementary failure, as eight knockout datasets with low target counts in individual layers lacked a *network-based* prediction (**Supplementary Figure 9A**), whereas eight knockout datasets missing data in at least one layer were predicted by the *network-based* strategy alone (**Supplementary Figure 9B**). In a second benchmark, GSEA showed that, on average, 25% of predicted GO terms were significantly enriched in the corresponding knockout datasets (GSEA, P < 0.05; **Fig 2E**). However, when we tested whether this recovery exceeded background enrichment in each knockout transcriptome using size-matched random GO term sets, the predicted sets did not outperform these random controls. This likely reflects context mismatch, as only four of the 13 knocked-out TFs have tissue-restricted expression (tissue-specificity index τ; Kryuchkova-Mostacci and Robinson-Rechavi, 2017; **Supplementary Figure 9C,D**), so single-condition knockouts of broadly expressed TFs capture only a fraction of their regulatory roles. Together, these knockout benchmarks show that network-based integration tolerates missing layers and outperforms overlap strategies where both yield predictions, while remaining sensitive to low target counts and context mismatches in single-condition knockouts.

Finally, to evaluate false-positive rates and confirm that the predicted GO terms capture genuine biological signal, we tested all three strategies on 3,000 degree-preserved randomized networks per TF, where lower enrichment indicates fewer false positives and higher specificity (**Supplementary File 2**). Across the 32 TFs annotated by all three strategies, the *network-based* method identified significant GO term enrichment in only ∼12% of randomized networks, compared with ∼28% and ∼72% for the *common-function* and *common-interaction* strategies, respectively (**Supplementary Figure 10A**). Relative to both overlap strategies, *network-based* integration also produced fewer significant GO terms per randomized TF’s targets, weaker significance (-log10 FDR), and lower semantic similarity to the GO terms predicted from the real network (**Supplementary Figure 10B–D**, **Supplementary Figure 11**). Conversely, when applied to the observed TF–target interactions, the *network-based* method recovered significantly more GO terms than expected by chance for 30 of the 32 TFs (**Supplementary Figure 12**). Therefore, we used *network-based* predictions exclusively hereafter.

### Prioritization of regulators by biological process

A major goal of multi-omic integration is to prioritize specific TF functions that guide both hypothesis- and data-driven experimental design. Our network-based strategy yielded 7,722 TF–GO associations (**Fig 2A** and **2C; Supplementary Figure 6B and 6C**), reflecting the combinatorial nature of transcriptional regulation in which individual TFs regulate multiple processes and a process may be regulated by multiple TFs. To narrow down this broad catalog into tractable specific processes, we restricted these associations to GO biological-process terms and collapsed terms with fewer than 50 annotated genes into their nearest parent, leaving 4,337 associations across 902 TFs and 559 terms in total (**Fig 3A** and **3B**). Term size explained little of the variation in the number of TFs per term (R² = 0.14, **Fig 3C**), so terms are not favored simply for being broadly annotated. TFs with larger neighborhoods, however, were associated with more terms (R² = 0.27, **Fig 3D**). Because degree is therefore not directly comparable across TFs, we computed a scaled enrichment score (Z-score) per TF and per GO term (*Methods*). Only 3.5% of associations (151/4,337) scored high (Z ≥ 1) in both directions (**Fig 3E**), indicating that most TFs and most terms carry several associations of comparable strength. Combining these two Z-scores into a reciprocal Z-score (rZ; *Methods*) provides a single specificity metric that highlights associations that are simultaneously unusually strong for the TF and unusually strong for the biological process, which we use for ranking below.

**Fig 3.**
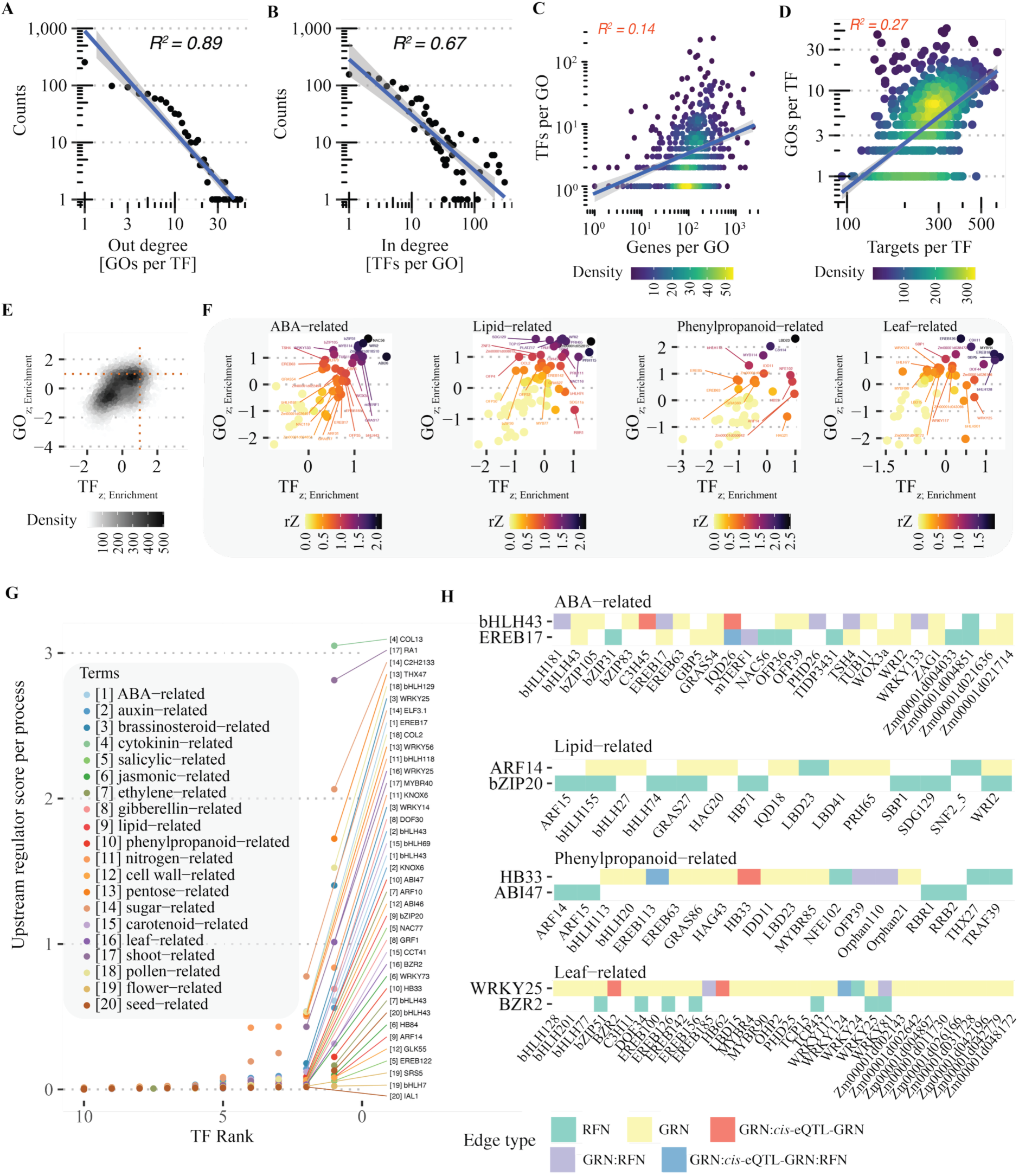
Prioritization of regulators by biological process using network-based predictions. (A, B) Out-degree (A) and in-degree (B) distributions of the TF–GO term predictions. (C, D) Density scatterplots of the number of TFs per GO term (C) and GO terms per TF (D) versus genes annotated per GO term and target genes per TF, respectively. (E) Density scatterplot of scaled TF–GO enrichment scores (GO_z and TF_z); orange dashed line marks associations with Z ≥ 1 for both. (F) Reciprocal Z-scores (rZ) for four processes (ABA-, lipid-, phenylpropanoid-, and leaf-related) mapped onto the scaled-score coordinates of (E); labels for TFs with rZ ≥ 0.5. (G) TFs ranked by upstream regulator score per process (labels for rank ≤ 2; bracketed indices match the process labels). (H) Heatmap of the top two upstream-scoring TFs (y-axis) per process and their targets (x-axis), colored by source network(s).

Grouping GO terms into higher-level processes revealed several strongly associated TF–process pairs (**Fig 3F, Supplementary Figure 13**; maximum Z-score in both directions). Several processes stood out, including abscisic acid (ABA) metabolism (46 associations, 44 TFs), lipid metabolism (62 associations, 55 TFs), phenylpropanoid metabolism (47 associations, 47 TFs), and leaf development (50 associations, 50 TFs). Filtering at rZ ≥ 0.5 narrowed these to 27, 25, 15, and 19 candidate regulators, respectively (**Fig 3F**). Among them, NAC56 and WRINKLED2 (WRI2) were associated with ABA, and WRI2 also ranked in the top three for lipids. Thirteen of the 47 phenylpropanoid-associated TFs, including OFP39, EREB149, WOX9a, HAG43, and LBD24, had been identified as regulators of phenolic-gene promoters in yeast one-hybrid screens (Yang *et al*. 2017; Abnave *et al*. 2024). Consistent with this overlap, our combined network links TFs to the 75 phenolic-gene promoters assayed in those screens 1.7 times more often than degree-matched random networks (permutation P < 10⁻⁴), driven by TF–biosynthetic gene interactions in the expression layer. Together, reciprocal Z-score prioritization narrows thousands of multifunctional associations to compact, process-specific shortlists of candidate regulators.

Biological regulatory networks are often organized hierarchically through TF–TF cascades and feed-forward loops that reinforce control and concentrate coordination in upper-level regulators (Alon 2007; Brkljacic and Grotewold 2017; Abnave et al. 2024). Here, to identify such higher-level regulators, we defined an upstream regulator score, the fraction of a TF’s within-process predicted targets that are themselves TFs. A high upstream regulator score indicates that a TF sits above other regulators in the same process, a proxy for process-level feed-forward loops (**Fig 3G**). Across 20 processes (eight hormone-related, seven metabolic, five developmental), cytokinin-related terms scored highest, led by COL13, followed by shoot-related terms, led by RA1. COL13 has been linked to carbon metabolism (Tu *et al*. 2020) and is differentially expressed in *id1* mutants (Minow *et al*. 2018), suggesting a role connecting cytokinin, carbon metabolism, and flowering (Bartrina *et al*. 2011; Wahl *et al*. 2013). Similarly, RA1 recovers its documented role in shoot development (Eveland *et al*. 2014), supporting the score. Additionally, tracing the top two upstream-scoring TFs per process across the individual network layers (**Fig 3H**) showed that each upstream regulator directly targets a top-rZ TF, including bHLH43 targeting EREB17 and WRI2 in ABA metabolism (with EREB17 also targeting WRI2 and NAC56 in a feed-forward loop), ARF14 targeting WRI2 and PRH65 in lipid metabolism, HB33 targeting LBD23 in phenylpropanoid metabolism, and WRKY25 targeting MYBR4, EREB126, and BZR2 in leaf development. Thus, these provide specific interaction candidates for experimental validation.

### Mapping TF–process associations to specific conditions

We previously showed that most TFs carry multiple predicted functions, and only 3.5% of associations stand out with high specificity in both directions (**Fig 3E**), leaving it unresolved which tissues, genotypes, or stress conditions drive each TF–process link. To further characterize these associations, we mapped the 4,337 TF–GO associations onto specific conditions using GSEA across individual expression datasets (*Methods*), namely the 45 datasets underlying the RFNs (Zhou et al. 2020) plus six heat- and cold-stress datasets (Zhou et al. 2022). Rather than using correlation to discover new associations, we ranked all genes within each dataset by their Pearson correlation with each TF (Gomez-Cano et al. 2022) to score where existing predictions are active. In total, 76% of tested TF–GO terms showed enrichment (FDR ≤ 0.1) in at least one dataset (**Supplementary Figure 14A-C**). Sorting the tested TFs into terciles by the mean percentage of their predicted GO terms enriched across datasets defined high (277 TFs, 72.5% enriched), intermediate (276 TFs, 53.3%), and low (276 TFs, 34.0%) tiers (Fig 4A, Supplementary Figure 14D). Low-tier TFs were enriched in fewer datasets (Supplementary Figure 14E) despite carrying more GO terms than high-tier TFs (Supplementary Figure 14F, P < 0.001, Wilcoxon), indicating that their predictions are more condition-restricted.

**Fig 4.**
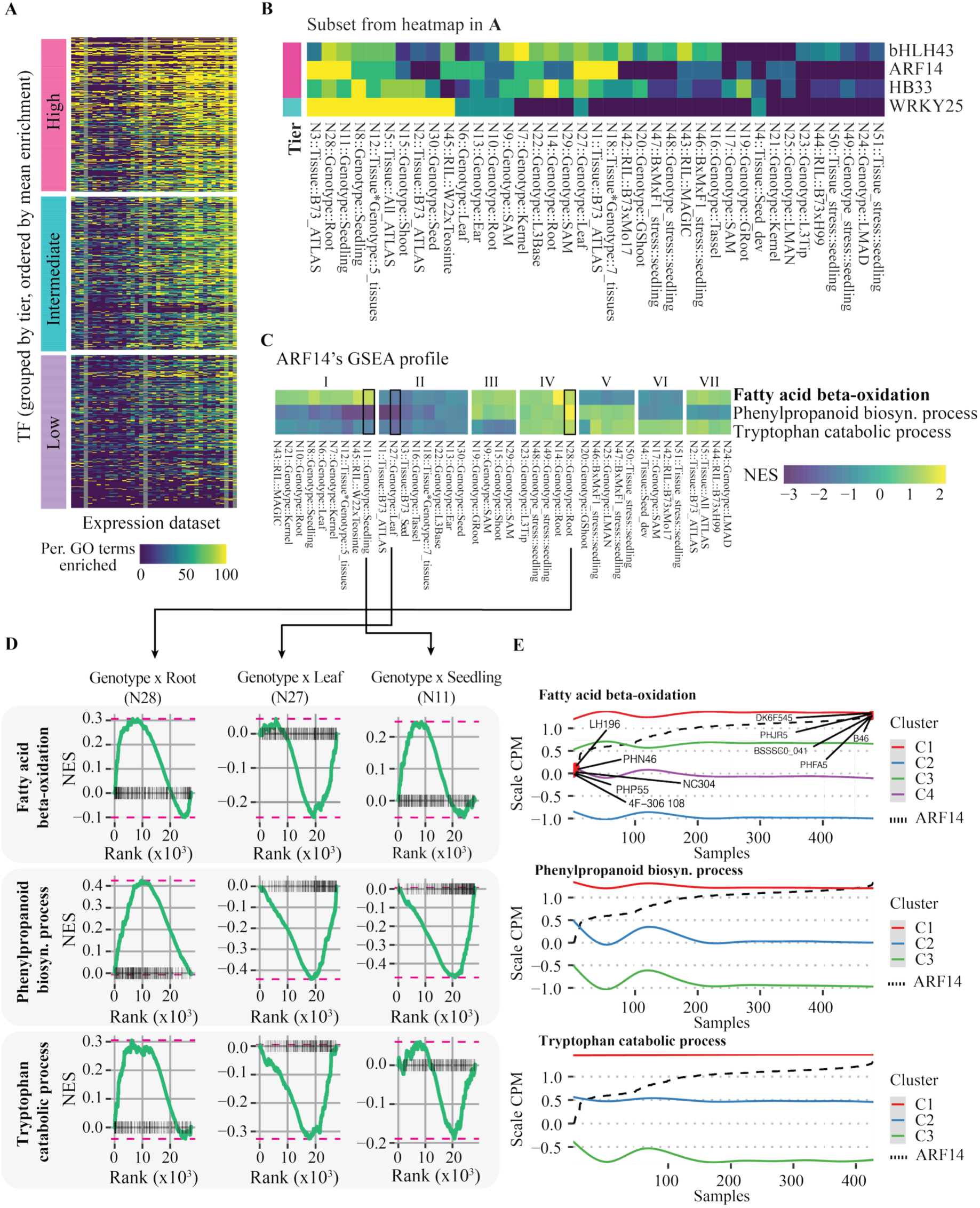
Mapping TF–GO term associations to specific conditions. (A, B) Heatmaps of the percentage of GO terms significantly enriched (GSEA) per TF (y-axis) across expression datasets (x-axis), for all TFs (A) and four example TFs (B). In (A), TFs are grouped into three deterministic tiers (high, intermediate, low) defined as terciles of the mean percentage of enriched GO terms. (C) Normalized enrichment score of ARF14 and its three associated GO terms across 39 datasets (absent in 12 datasets lacking ARF14 expression or significant enrichment). (D) GSEA profiles for three datasets from (C), showing gene-set accumulation along the ARF14-correlation ranking. (E) LOESS plot of scaled expression levels for the gene sets in (D) across samples ordered by ARF14 expression (dashed line, scaled ARF14 expression); genes clustered after normalization.

Focusing on the upstream regulators (defined in the previous section) bHLH43, ARF14, and HB33 (high tier) and WRKY25 (intermediate tier), we found significant TF–GO enrichment across nearly all datasets (Fig 4B), supporting their predicted upstream roles. The GSEA normalized enrichment score also revealed condition- and process-specific activity (**Fig 4C**, **Supplementary Figure 15**). ARF14, for example, was predicted as a lipid regulator through fatty-acid β-oxidation (GO:0006635) and was additionally enriched for tryptophan catabolism (GO:0006569), and phenylpropanoid biosynthesis (GO:0009699), three interdependent pathways that share metabolic precursors (Maeda and Dudareva 2012; Perez de Souza et al. 2020) and whose enrichment scores reverse sign across conditions (illustrated by datasets N28, N27, and N11; **Fig 4C, 4D**). This sign shift suggests condition-dependent positive or negative regulation across subsets of genes within each pathway. Notably, one of these datasets (N11) captures expression variation across diverse inbred lines, separating an ARF14-independent cluster (C1) and pinpointing lines with extreme ARF14 expression where perturbing ARF14 should most strongly affect its target pathways (**Fig 4E**).

### Topological properties predict TF paralog redundancy

Predicting functional redundancy among maize paralogs remains difficult (Schnable *et al*. 2011; Li *et al*. 2016; Kono *et al*. 2018; Han *et al*. 2023). We reasoned that functionally redundant paralogs would share topological properties in the integrated network. To test this hypothesis, we quantified pairwise embedding similarity across all genes in the network (**Fig 5A**, *Methods*). For 932 maize TF paralog pairs present in our datasets (Zhang *et al*. 2017), we compared both the direct embedding similarity between the two paralogs and the correlation of their genome-wide similarity profiles across all other genes (**Fig 5A**). Paralogs on the same chromosome (a proxy for tandem or local duplicates) showed higher scores than paralogs on different chromosomes by both metrics (**Fig 5B, 5C**). Combining both scaled metrics (correlated, Pearson r = 0.68) identified highly similar pairs (**Fig 5D**). At the extreme scores, the ten most-similar paralog pairs shared many network-associated genes, whereas the ten least-similar pairs shared none (**Fig 5E**). Grouping all pairs into nine similarity bins from lowest (bin I) to highest (bin IX) across both metrics (**Fig 5D, 5F**) showed that bin IX contained 5–7-fold more same-chromosome paralog pairs than the other bins (**Fig 5F**). Consistently, comparing shared network associations (Jaccard index of each pair’s associated genes) against a null distribution of neighborhood-size-matched random TF pairs (*Methods*) showed that only bin IX exceeded the null expectation (3.6-fold higher than expected, empirical P < 0.001, **Fig 5G**), whereas the least-similar pairs (bin I) shared fewer associations than expected by chance (**Fig 5G**). The large sharing score in bin IX therefore reflects genuine topological similarity rather than the shared origin of the annotations.

**Fig 5.**
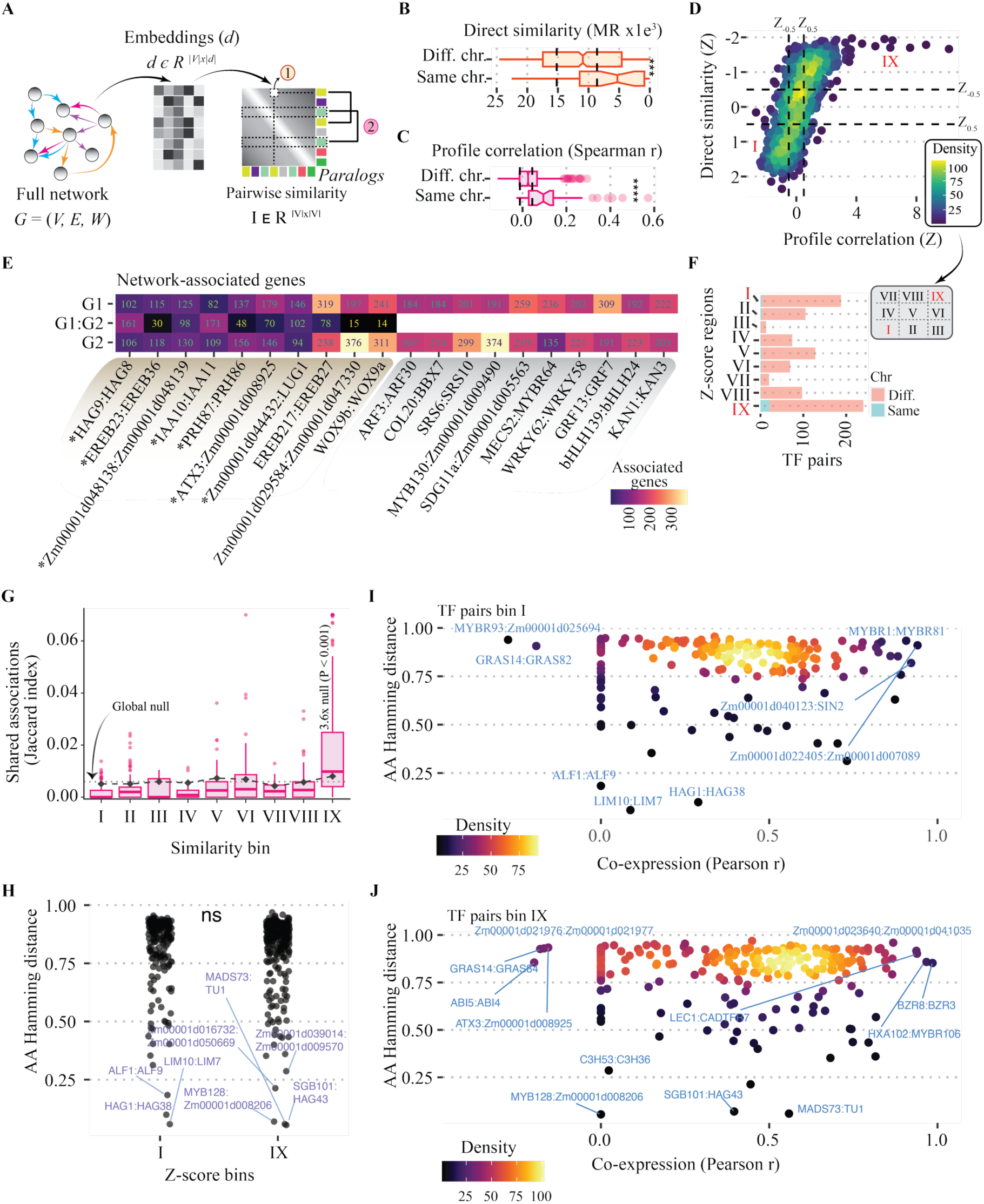
Network embeddings predict TF paralogs with functional variation. (A) Diagram of the paralog-comparison steps using embedding similarities. (B, C) Direct embedding similarity of TF paralog pairs (B) and Spearman correlations of their genome-wide similarity profiles (C). (D) Combined scaled embedding-similarity and profile-correlation scores per TF pair (dashed lines at Z = ±0.5). (E) Heatmap of associated-gene counts for the top-ten most-and least-similar TF pairs (G1, G2 = genes unique to each TF; G1:G2 = shared). (F) TF pairs per similarity bin (I–IX, mapped from D). (G) Jaccard index of shared associations for paralog pairs in each similarity bin (boxes). The dashed line marks the global null, calculated as the mean Jaccard of 20,000 random TF pairs. Black points mark the size-matched null for each bin, the mean Jaccard of random TF pairs drawn from the same neighborhood-size quintiles as the paralogs in that bin (20 per pair). (H) Amino-acid Hamming distance between TF pairs in bins I and IX. (I, J) Amino-acid Hamming distance versus co-expression (Pearson r) for bin I (I) and bin IX (J). Asterisks as in Fig 2.

We next asked whether low topological similarity reflects divergence in protein sequence or regulation. Comparing bins I and IX (the most different bins), we found no significant difference in amino-acid sequence (**Fig 5H**). However, mapping co-expression against sequence similarity distinguishes paralogs potentially undergoing neo- or subfunctionalization (**Fig 5I, 5J**), as the contrasts separated pairs whose protein sequences diverged despite shared expression from pairs that retained protein conservation while partitioning expression domains or regulatory targets. Examples include MYBR1/MYBR81 (distinct embeddings, amino-acid Hamming distance ∼0.9, and high co-expression, Pearson r > 0.9; **Fig 5I**), MADS73/TU1 (similar embeddings, amino-acid Hamming distance ∼0.05, and similar expression, Pearson r ∼0.55; **Fig 5J**), and C3H53/C3H36 (similar embeddings, amino-acid Hamming distance ∼0.3, but low co-expression, Pearson r ∼0.1; **Fig 5J**). Because sequence and co-expression comparisons are partly independent of network topology, combining all three metrics prioritizes specific paralog pairs as testable candidates for functional redundancy or divergence.

## Discussion

Cells rely on complex molecular networks to coordinate biological processes, and the rapid expansion of plant multi-omics datasets has increased the interest in methods for the accurate integration of diverse molecular measurements (Wen *et al*. 2016, 2018; Liu *et al*. 2016; Walley *et al*. 2016; Jin *et al*. 2017; de Abreu E Lima *et al*. 2018; Schaefer *et al*. 2018; Lee *et al*. 2019). Many established multi-omics integration methods assume that different data types are measured in the same samples (Wang *et al*. 2014; Argelaguet *et al*. 2018). Because plant expression atlases, PDI assays, and population-scale eQTL datasets are typically generated independently across genotypes, tissues, and TF subsets, we compared three gene-level integration strategies across four molecular network layers. Summarizing the combined network through gene embeddings outperformed target- or function-overlap strategies in recovering knockout GO terms, annotated over eightfold more TFs, and showed lower false-positive enrichment in randomized networks. Crucially, embeddings remained informative when individual layers were missing. This last, a practical necessity because in maize, as in many other crops and non-model organisms, PDI data cover only a small fraction of TFs. Thus, we envision this strategy proving useful for systems studies, combining whatever uneven molecular layers are available to enable functional inference.

Focusing on network-based predictions to prioritize regulators of specific processes yielded insight into maize metabolism and development. Only 3.5% of predictions had a single dominant TF per GO term, consistent with most TFs contributing to multiple functions and most processes requiring multiple TFs (Yang *et al*. 2017; Tu *et al*. 2020; Tang *et al*. 2021; Gray and Grotewold 2025). This is a hallmark of GRNs, whose regulatory capacity is amplified through TF–TF interactions (Reményi *et al*. 2004; Brkljacic and Grotewold 2017), as observed in maize at the TF–target (Yang *et al*. 2017; Tu *et al*. 2020) and *cis*-regulatory levels (Marand *et al*. 2021). We also predicted feed-forward motifs, a known signal-reinforcement mechanism (Alon 2007), extending the concept to the level of biological processes, with implications for targeted metabolic engineering. Because TFs act in a condition-specific manner (Sonawane *et al*. 2017), we mapped predictions across 51 expression datasets (Zhou *et al*. 2020, 2022) using GSEA, of which 39 returned enrichment results. Most TFs (the high and intermediate tiers) regulate multiple GO terms across conditions, while a subset (the low tier) act in few conditions, underscoring a condition-specific regulatory landscape shaped by the combined action of multiple TFs (Sonawane *et al*. 2017). ARF14 illustrates the findings. This auxin response factor (Cancé *et al*. 2022) is predicted to regulate several metabolic and developmental processes, including tryptophan catabolism. Since tryptophan is a precursor of the auxin indole-3-acetic acid (Zhao 2010), this suggests a mechanistic link between hormonal and metabolic control, with condition-specific coordination between ARF14 and each pathway (Vanstraelen and Benková 2012; Derksen *et al*. 2013; Munné-Bosch and Müller 2013; Pacifici *et al*. 2015; Sawicki *et al*. 2015).

Functional redundancy between maize subgenomes remains incompletely understood (Schnable *et al*. 2011; Li *et al*. 2016; Kono *et al*. 2018; Han *et al*. 2023). Using embedding similarity, we found that TF paralogs with more similar embeddings share more targets and are more often tandem duplicates (**Fig 5E–G**), with sharing in the top-similarity bin exceeding a size-matched random expectation. Although amino-acid divergence did not differ between high- and low-similarity paralogs (**Fig 5H**), co-expression was slightly higher among topologically similar pairs. Integrating protein sequence, expression, and embedding similarity identifies specific candidates for neo- or subfunctionalization and partial redundancy (**Fig 5I, 5J**) for experimental testing.

Overall, integrating multi-omic datasets through embeddings followed by clustering identifies genes with similar wiring in the fully integrated network. This agrees with earlier multi-network embeddings in yeast and human, where compact representations of combined topology predicted gene function better than any single network (Cho *et al*. 2016). Those methods, and more recent ones such as BIONIC (Forster *et al*. 2022), were developed and benchmarked in yeast and human, where thousands of mutant phenotypes and curated annotations are available for evaluation. Our results show that this principle transfers to crops despite sparser, unevenly distributed data and few characterized mutants. Because embedding similarity connects > 24K maize genes, the same framework can annotate other gene classes or pathways and offers a general template for integrating unmatched functional genomics datasets in non-model species.

## Limitations

Several limitations should be considered when interpreting these results. First, RFN edges derive from random-forest regression of expression data and represent predictive rather than causal associations. Second, annotations depend on the neighborhood cutoff D; although overall yield was stable across a 40-fold threshold range (**Supplementary File 3, Supplementary Figure 16**), boundary associations varied, so we provide per-association stability scores (**Supplementary Table 8**) to prioritize threshold-robust predictions. Third, modest agreement with knockout transcriptomes likely reflects condition mismatches and the small benchmark size (14 maize TFs), which could be expanded in species with larger mutant collections such as Arabidopsis thaliana. Fourth, mapping to the B73 AGPv4 reference omits pangenome variation absent from B73. Fifth, the networks do not distinguish activating from repressive regulation. Sixth, layer weights were set by support count rather than fitted; approaches that tune data-type weights against prediction error (Cassan *et al*. 2024) could potentially improve both layer weighting and construction of the random-forest network. Finally, we did not compare the embedding method used here, node2vec as implemented in PecanPy (Liu and Krishnan 2021), with alternative multi-network embedding approaches such as Mashup (Cho *et al*. 2016) or BIONIC (Forster *et al*. 2022). Our analysis therefore evaluates strategies for combining network layers rather than the relative performance of embedding algorithms, and alternative embedding methods could produce different results.

## Materials and Methods

### Identification of eQTL

A set of 304 diverse inbred lines with publicly available SNP and gene-expression data was included in the eQTL analysis (Kremling *et al*. 2018; Bukowski *et al*. 2018; Mazaheri *et al*. 2019). Because the whole-genome-sequencing SNPs and the RNA-seq-derived markers were both mapped to the B73 maize reference genome (AGPv4) (Jiao *et al*. 2017), we concatenated the datasets with bcftools (v1.7) (Danecek *et al*. 2021), preferentially keeping the RNA-seq marker where the two overlapped. The expression datasets capture both genotypic and tissue-level variation.

### Expression data and random-forest regulatory networks (RFNs)

All expression and RFN datasets used here are listed in **Supplementary Table 3** and were previously published (Zhou *et al*. 2020), except RFN 46, which we built from the 304 inbred lines analyzed for genetic markers (n304). We gathered pre-computed counts-per-million (CPM) values for these lines and applied the exact procedure of Zhou *et al*. (2020). All RFNs were inferred with RandomForestRegressor, retaining the top 100K associations per network. These networks predict regulatory (TF to target) relationships from expression and do not, by themselves, imply correlation-based co-expression or causal regulation.

### eQTL identification and classification

eQTLs were identified across eight tissues spanning germination to maturity (GRoot, Gshoot, Kern, L3Base, L3Tip, LMAD, LMAN, seedling) (Kremling *et al*. 2018; Bukowski *et al*. 2018; Mazaheri *et al*. 2019). SNPs were filtered to biallelic markers with minor-allele frequency ≥ 0.05, and expression sets to genes with ≥ 6 reads and ≥ 0.1 TPM in ≥ 20% of samples, leaving 15.5–16.7M SNPs tested against 15.3K–26.4K genes per tissue.

Each SNP–gene association was evaluated with eight candidate linear models fitted in rMVP (Yin *et al*. 2021) and run as an HTCondor array (R 4.0.2). The first was a naive single-marker model with no covariates (method “GLM”). The remaining seven were mixed linear models (method “MLM”, variance components by Haseman–Elston regression) that fitted a marker-based genomic relationship matrix as a random polygenic effect and differed only in their fixed-effect covariates, which were none, the first five principal components (PCs) of the SNP matrix, or those five PCs plus the top 5, 10, 15, 20, or 25 PEER factors (Stegle *et al*. 2012) of the tissue expression matrix. The relationship matrix and SNP PCs came from the filtered marker set via MVP.Data, and 60 PEER factors per tissue were estimated with the GTEx v8 eQTL pipeline, of which the top 25 were retained.

Significance thresholds came from covariate-free permutations in tensorQTL (Taylor-Weiner *et al*. 2019) (10,000 permutations), applied as a per-gene nominal P-value threshold for cis associations (±100 kb window) and a single per-tissue threshold for trans associations, set at the fifth percentile of the permuted minimum P-value distribution. For each SNP–gene pair we recorded how many of the eight models passed the applicable threshold. Cis and trans associations were then filtered differently, with trans associations retained for the *trans*-eQTL network requiring support from at least two models and the cis-eQTL set used for *cis*-eQTL-supported GRN construction requiring support from all eight. Associations involving non-syntenic target genes (Zhang *et al*. 2017) were discarded, and the remainder were classified as *cis*-eQTLt, *trans*/*cis*-eQTL, *cis*-eQTL, *trans*-eQTL, or unassigned, based on the distance between the eQTL and its target and co-location with annotated genes (B73 v4, **Supplementary Figure 2A**).

### Protein–DNA interaction (PDI) data analysis

Raw reads from plant ChIP-seq (Bolduc *et al*. 2012; Morohashi *et al*. 2012; Eveland *et al*. 2014; Pautler *et al*. 2015; Zhan *et al*. 2018; Li *et al*. 2018; Dong *et al*. 2019), protoplast ChIP-seq (pChIP-seq) (Tu *et al*. 2020), and DAP-seq (Ricci *et al*. 2019; Dong *et al*. 2020) were collected from public datasets (Supplementary Table 2). Read QC and peak calling followed (Gomez-Cano *et al*. 2022). Briefly, read quality was assessed with FastQC (v0.11.5). Adapters and low-quality reads were trimmed with Trimmomatic (Bolger *et al*. 2014) (ILLUMINACLIP:Adapter.fastq:2:40:15 SLIDINGWINDOW:4:20 MINLEN:27). Reads were mapped to B73 AGPv4 (Jiao *et al*. 2017) with Bowtie2 v2.3.5.1 (Langmead and Salzberg 2012) (nuclear chromosomes only). Alignments with MAPQ < 30 were removed with Samtools v1.9 (Li *et al*. 2009), and duplicate reads were removed with Picard MarkDuplicates. Peaks were called with GEM v3.4 (Guo *et al*. 2012). For DAP-seq, HALO-only sequences served as negative control. Plant ChIP-seq was called with replicates using the corresponding mutant or tagged-protein control, and pChIP-seq was called with replicates. All runs used the default read distribution with --k_min 6 --k_max 15 –relax --outNP --sl. Only TFs with > 500 peaks were retained. *Peak quality control.* TF peaks were kept if they overlapped ACRs and had a per-assay CPM Z-score > −0.5. CPMs per peak were obtained by extending each summit 50 bp per side, converting to SAF, counting reads with Rsubread v1.32.2 (Liao *et al*. 2019), and CPM-normalizing. ACRs were collected from (Marand *et al*. 2021). *Peak–TSS distance thresholds.* The proximal cutoff (≤ 3 kb) captures promoter-proximal binding and matches the concentration of retained high-coverage peaks at ACRs near the TSS (**Supplementary Figure 3D and 3E**). The distal window (> 3 kb to ≤ 50 kb) spans the range within which most characterized maize distal *cis*-regulatory elements and distal ACRs occur (Ricci *et al*. 2019; Marand *et al*. 2021). Distal peaks were retained only with independent cis-eQTL support (peak summit within 20 bp of a cis-eQTL), which limits spurious long-range target assignments.

### Functional annotation

Functional annotations used only sorghum-syntenic genes (Zhang *et al*. 2017). PWYs were from CornCYC (Andorf *et al*. 2016) and GO terms from GAMER (Wimalanathan *et al*. 2018). Syntenic genes followed Zhang et al. (2017). PWY and GO enrichment used the R packages GeneOverlap (v1.30.0) and topGO (v2.46.0), respectively. GO semantic similarity used GOSemSim (v2.20.0) with the “Wang” method (Yu 2020). For common-function analyses, GO comparisons used the original enriched-term sets. Afterward, GO terms were mapped to their nearest parent with Rrvgo (v1.6) (Sayols 2023) to reduce redundancy, as were significant terms from common-target and network-based analyses.

### Network integration

#### Common interactions

We counted how often each TF–target association appeared across the four layers (trans-eQTL network, GRN, cis-eQTL-supported GRN, and RFNs), retaining as “common” those present in ≥ 2 layers for the same TF. We then tested PWY/GO enrichment among each TF’s common targets and mapped significant GO terms to nearest parents with Rrvgo (Sayols 2023). *Common function.* For each TF and layer, we tested PWY/GO enrichment among targets. TFs with ≥ 1 term enriched in ≥ 2 layers were retained (Supplementary Figure 5A). PWY similarity across layers was assessed per TF by Fisher’s exact test (a gene can map to multiple PWYs). GO similarity used semantic similarity, followed by mapping significant terms to nearest parents. *Network-based.* All four layers were combined and interaction frequencies scaled from 0.5 to 1 before embedding. Embeddings were computed with PecanPy (Liu and Krishnan 2021) (--weighted --dimensions 50 --walk-length 80 --num-walks 10 --directed). Gene similarity was the mutual rank of mutual information, MRMI = sqrt(MI_rank × tMI_rank), where MI (Parmigene (Sales and Romualdi 2011)) is ranked and tMI_rank is the transposed rank matrix. Genes highly similar to a TF (its MRMI neighborhood) were selected with a decay function D = e^(−(MRMI − 1)/50), keeping genes with D ≥ 0.005 (MRMI ≤ ∼266), then tested for PWY/GO enrichment (significant terms mapped to nearest parents). The combined network is a directed, weighted edge list connecting 36.4K genes through 4.6M unique interactions (interactions deduplicated within each layer before combining). Each interaction was weighted by the number of layers supporting it, w = 0.5 + 0.125 × n with n = 1–4, giving weights from 0.625 (single layer) to 1.0 (all four layers). This scaling lets interactions with multi-layer support guide the random walks more strongly (up to 1.6-fold) while keeping single-layer interactions visitable, so no layer is excluded from the embedding.

#### Threshold-sensitivity analysis

To assess the dependence of network-based annotations on the neighborhood cutoff, we repeated neighborhood definition and PWY/GO enrichment at five thresholds, D ≥ 0.005 (the value used throughout), 0.02, 0.05, 0.10, and 0.20, corresponding to MRMI ≤ 266, 197, 151, 116, and 81. At each threshold we recomputed the enrichment background from the thresholded network and applied the same tests and multiple-testing corrections as the main analysis. Robustness was quantified as (i) the number of annotated TFs and significant associations per threshold, (ii) retention of baseline associations and Jaccard similarity of significant sets, (iii) the rank correlation of enrichment P-values against the baseline, and (iv) a per-association stability score, defined as the number of thresholds (out of five) at which the association was significant, reported in Supplementary Table 8. For TF–GO associations, stability was assigned at the level of the released parent terms through the recorded term-to-parent mapping. Associations whose terms are obsolete or restructured in the current GO release are flagged as not assessable rather than scored.

#### Trans-eQTL network class-frequency normalization

To test whether gene-class contributions to the *trans*-eQTL network reflect class SNP frequency, we counted, per class (TF, co-regulators, Mediator, kinases, enzymes, other), the number of class genes, unique trans-eQTL SNPs co-locating with class genes, source genes, interactions, and unique targets. Normalized rates were computed as interactions per SNP and per class gene. Genes annotated to multiple classes were assigned by the priority TF > co-regulators > Mediator > kinases > enzyme.

#### Paralog null model

To test whether shared network associations among similar paralogs exceed chance expectation, we compared each similarity bin against random TF pairs. For each of the 932 paralog pairs we sampled 20 random non-paralog TF pairs matched on neighborhood size (quintiles of the MRMI-neighborhood size distribution across the 2,915 TFs with neighborhoods, fixed seed). Per-bin empirical P-values were obtained by bootstrap of the matched-null bin means (B = 2,000). A global null of 20,000 unmatched random TF pairs gave consistent results.

#### Evaluation of functional predictions

Predicted PWYs/GO terms were compared with those enriched among DEGs (and their log2 fold-changes) in loss-of-function mutants. Knockout data were from KN1 (Bolduc *et al*. 2012), RA1 (Eveland *et al*. 2014), FEA4 (Pautler *et al*. 2015), O2 (Zhan *et al*. 2018), bZIP22 (Li *et al*. 2018), and TB1 (Dong *et al*. 2019), plus MYBR32_m1, WRKY82_m1, HSF13m1m2, HSF18m1, HSF20m1, HSF29m1, HSF29m2, WRKY2m2, WRKY8m1, and WRKY8m2 (Ellison *et al*. 2023) (DEGs listed in **Supplementary Table 9**). DEGs were selected by DESeq2-adjusted P < 0.05 (Love *et al*. 2014) and enriched as above. Prediction– observation similarity used PWY overlap and GO semantic similarity. GSEA used FGSEA (v1.18) (Korotkevich *et al*. 2016) (minSize = 5, maxSize = 1000, eps = 0), with gene sets defined by each TF’s predicted PWYs/GO terms. The recovered fraction was the number of significant (P < 0.05) terms over the number tested. For random-network comparisons, each of the four networks was randomized 3,000 times (“rewire”, R igraph v1.2.4.1, no loops, niter = nodes × 10,000), then annotated and integrated as above.

#### Prioritization of transcriptional regulator–process associations

All prioritization used network-based results. GO terms with < 800 genes were retained, and overly specific terms (< 50 genes) were mapped to their parent (Functional annotation). For each TF_i–GO_j association we computed an enrichment score E_ij = log2[(c/t)/(p/u)], where c is the overlap of TF_i targets and GO_j genes, t the TF_i target count, p the GO_j gene count, and u the total syntenic-with-sorghum gene count (Zhang *et al*. 2017). E_ij was normalized per TF and per GO: Z_i = (E_ij − U_i)/σ_i and Z_j = (E_ij − U_j)/σ_j, where U and σ are the mean and SD of E over, respectively, all GO_j for TF_i and all TF_i for GO_j. The reciprocal Z-score was rZ_ij = sqrt(max(0, Z_i)² + max(0, Z_j)²). The upstream regulator score for a TF within a process was the number of its within-process targets that are themselves TFs divided by its total TF targets, approximating the proportion of process-level feed-forward loops.

#### Determining sequence similarities among TF paralogs

Transcript sequences for each paralog pair were obtained from MaizeGDB (genome v4) (Jiao *et al*. 2017). TF similarity was the average Hamming distance across all transcripts of the two TFs, computed with DECIPHER (v2.22) (Wright 2016) (“DistanceMatrix”, includeTerminalGaps = TRUE, penalizeGapLetterMatches = TRUE, correction = “none”).

#### Mapping TF–GO associations to conditions by GSEA

GSEA followed (Gomez-Cano *et al*. 2022), using FGSEA v1.18.0 (Korotkevich *et al*. 2016) with Pearson correlation coefficients as the ranking metric. The weighted PCC (wPCC) between each TF and the genes annotated to each GO term was computed on log2-CPM values with wCorr (v1.9.1) (Emad and Bailey 2017) (threshold 0.4). Expression datasets and conditions were as defined above and in Zhou et al. (2022). GO gene sets were from GAMER (Wimalanathan *et al*. 2018). Of the 51 datasets tested, 39 returned GSEA results and constitute the columns of Fig 4A. TF tiers in Fig 4A were defined as terciles of each TF’s mean percentage of enriched GO terms across tested datasets, a deterministic grouping (cutoffs 45.5% and 60.9%) that replaces the original k-means clustering, for which standard diagnostics did not support a single k (**Supplementary Figure 16E–G, Supplementary File 3**).

## Use of generative AI

The authors wrote all the main text. Claude (Anthropic; models Opus 5.1 and Fable 5.1 through Claude Code) was used to edit and annotate code. No results or references were generated by the model, and the authors reviewed all output and took full responsibility for the content.

## Data Availability

The processed networks and predictions generated in this study (the four network-layer edge lists, the combined weighted network, gene embeddings, the TF to function catalog in Supplementary Table 8, and the prioritization scores) are deposited at Zenodo under DOI 10.5281/zenodo.21866340. All primary datasets reanalyzed here are publicly available: ChIP- and DAP-seq accessions are listed in Supplementary Table 2, expression data are from (Zhou *et al*. 2020, 2022), SNP data from (Kremling *et al*. 2018; Bukowski *et al*. 2018; Mazaheri *et al*. 2019; Yang *et al*. 2022), and ACRs from GEO accession GSE155178 (Marand *et al*. 2021). Analysis code is available at https://github.com/gomezcan/Multi-Omic_Network_Integration_Maize and archived at Zenodo under DOI 10.5281/zenodo.21875975 (release v1.0.0, MIT license).

## Author Contributions

**Conceptualization:** F.G.-C., J.R., P.Z.; **Methodology:** F.G.-C., J.R., P.Z., A.K., Y.-H.C., E.L.E., L.G.-C.; **Formal analysis / Investigation:** F.G.-C. (with input from E.G., N.M.S., N.d.L., A.K.); **Writing – original draft:** F.G.-C., E.G.; **Writing – review & editing:** all authors; **Supervision:** E.G., N.M.S., N.d.L.; **Funding acquisition:** E.G., N.d.L., N.M.S.

## Supplementary Material

**Supplementary File 1.** Expanded descriptions of TF functional annotation by common interactions and common function.

**Supplementary File 2.** Evaluation of functional predictions against random networks.

**Supplementary File 3.** Threshold-sensitivity analysis of network-based annotation.

**Supplementary Table 1.** Inbred lines and tissues used in the eQTL analysis.

**Supplementary Table 2.** ChIP- and DAP-seq datasets reanalyzed, including the SRA accession, library layout, method, TF identity and family, sample, condition, tissue, and replicate of each dataset.

**Supplementary Table 3.** Datasets used to build the RFN layer.

**Supplementary Table 4.** Expression datasets used for eQTL identification.

**Supplementary Table 5.** Gene categories used to classify trans-eQTLs.

**Supplementary Table 6.** Trans-eQTL associations for HSF20.

**Supplementary Table 7.** Total PDIs (peaks) used to build the GRN. Provided through the Zenodo deposit (DOI 10.5281/zenodo.21866340) because it exceeds supplementary file size limits.

**Supplementary Table 8.** Annotated functions (GO/PWY) per TF and method, the core TF to function catalog, including per-association threshold-stability scores (n of 5 thresholds significant). All TF–PWY associations carry stability scores. TF–GO associations are scored when assessable under the current GO release (5,469 of 7,722); associations whose terms are now obsolete or restructured in the ontology are flagged NA. The full table is deposited at Zenodo (DOI 10.5281/zenodo.21866340).

**Supplementary Table 9.** Differentially expressed genes per published maize TF knockout dataset. The compendium holds 44 datasets for 29 TFs; the Used_in_evaluation column flags the 21 datasets for 14 TFs used to evaluate the functional annotations, and the Dataset_key sheet summarizes all 44 with their mutant allele, tissue, source and differentially expressed gene count.

**Supplementary Figure 1.**
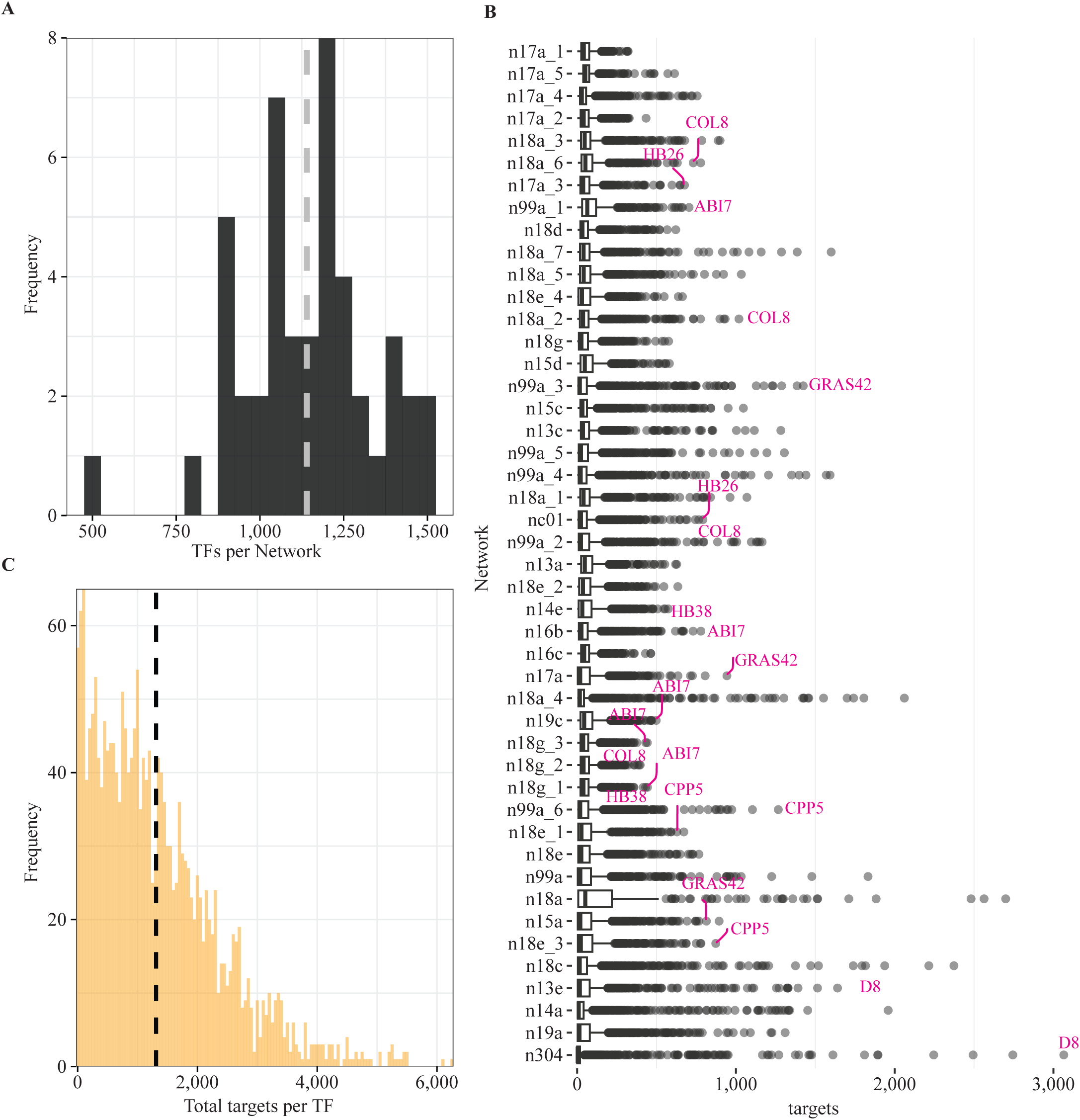
Total TFs and interactions in the random-forest regulatory network (RFN) layer. (A) Histogram of the number of TFs with at least one target gene per RFN. The dotted gray line indicates the average number of TFs across all 46 RFNs. (B) Boxplot of total target genes per TF across the RFNs. Magenta labels highlight TFs with the largest number of target genes in several RFNs. The RFNs shown in (B) are described in Supplementary Table 3. (C) Histogram of the total target genes per TF after combining the results of all 46 RFNs.

**Supplementary Figure 2.**
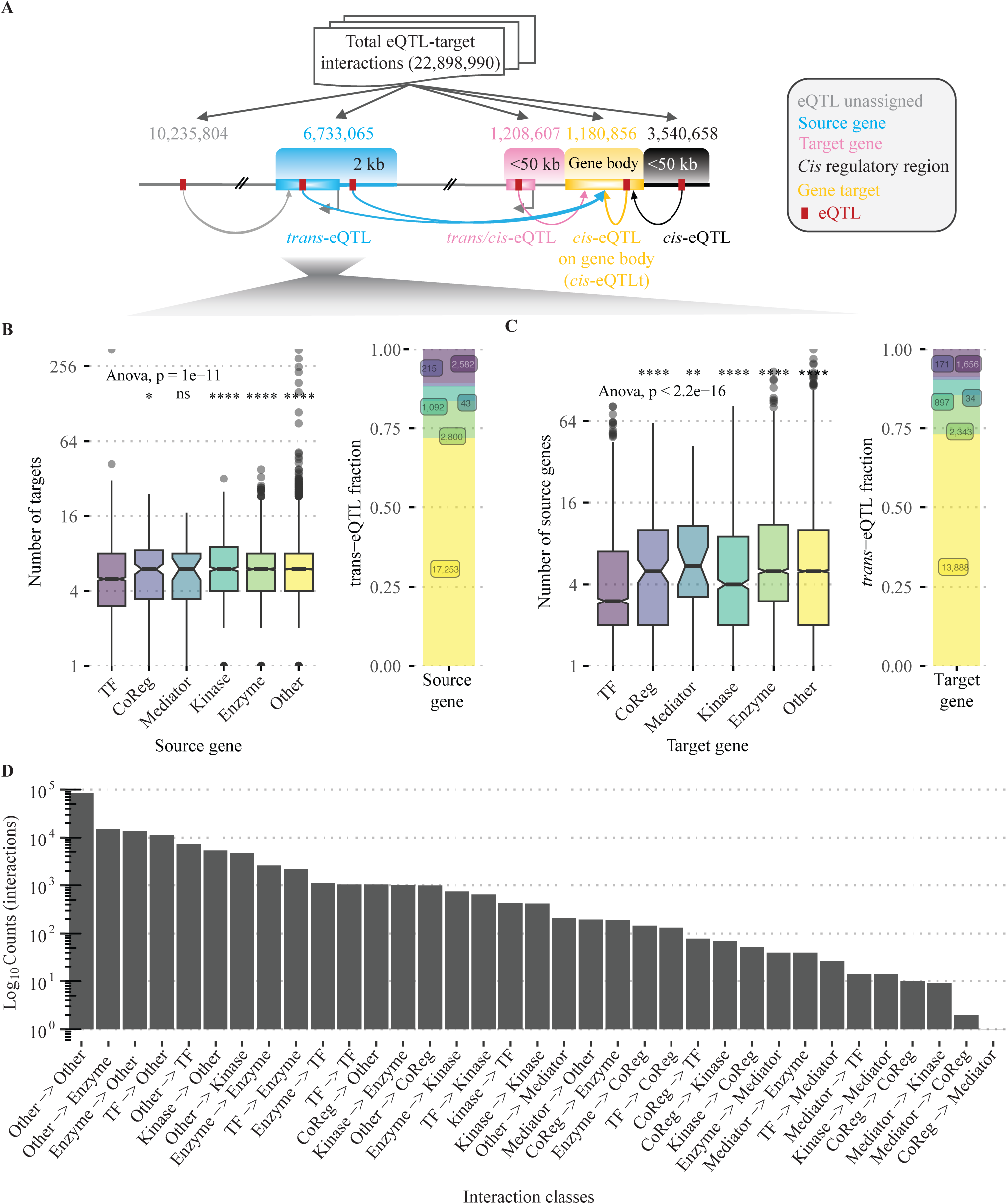
Defining the trans-eQTL network. (A) Model indicating the total eQTLs identified and the classification schema used to define trans-eQTL, trans/cis-eQTL, cis-eQTLt (eQTLs co-localized with their target genes), and cis-eQTL classes: 10.2M unassigned, 6.7M trans-eQTL, 1.20M trans/cis-eQTL, 1.18M cis-eQTLt, and 3.5M cis-eQTL. *Trans*-eQTLs were used to define the *trans*-eQTL network. A source gene (blue) is a gene whose promoter (2 kb upstream of the annotated TSS) or gene body overlaps an eQTL, and a target gene (yellow) is a gene whose expression is associated with that eQTL. (B, C) Classification of source (B) and target (C) genes into five functional categories (unclassified genes as other). Left panels show the number of genes per category, and right panels show the fraction of each category over all genes in the *trans*-eQTL network. (D) Total interactions by gene-category pair. Panels B– D show gene and interaction counts and fractions across functional classes (SNP-normalized interaction rates are reported in the text).

**Supplementary Figure 3.**
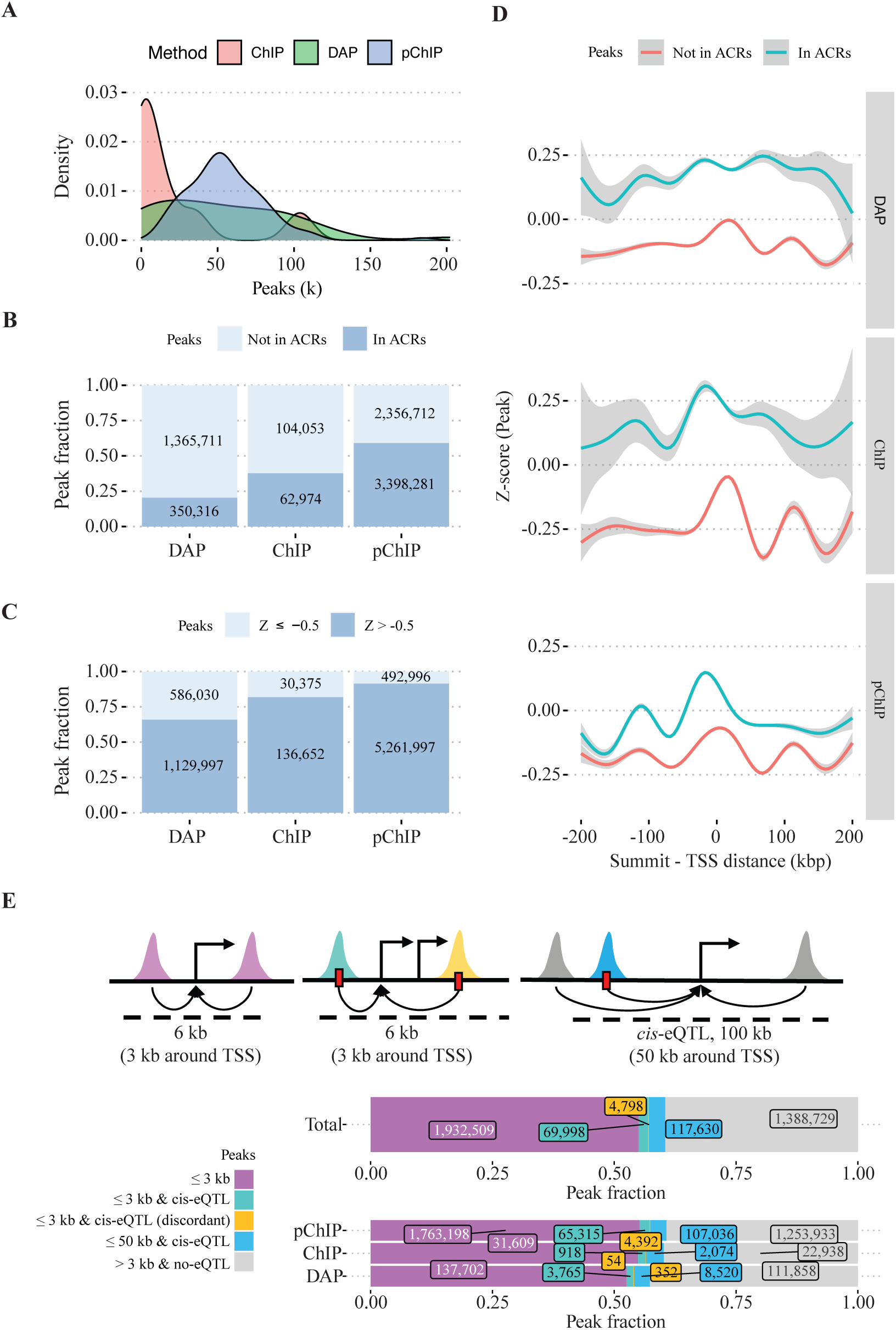
Establishing the PDI-based GRN layer. (A) Density plot of peak counts by PDI data type. (B, C) Fraction of peaks mapped to accessible chromatin regions (B) and fraction of peaks with low coverage (normalized read coverage Z ≤ −0.5) (C). Accessible-chromatin overlap filtering removed ∼3.8M peaks and coverage filtering removed a further ∼1.1M peaks, mostly from DAP-seq assays, leaving ∼3.4M high-confidence peaks. (D) LOESS plot of coverage Z-scores per peak in 10 kb bins within 200 kb of the closest transcription start site (TSS). Peaks with the highest coverage Z-scores mapped to accessible chromatin near the TSS across all data types. (E) Classification schema (top) and corresponding proportions of peaks (bottom) used to determine target genes. Peaks proximal to the TSS (≤ 3 kb) comprise peaks without cis-eQTL support (54.9% of analyzed peaks), peaks with cis-eQTL support and consistent target prediction (1.9%), and peaks with cis-eQTL support but discordant prediction (0.1%). Distal peaks comprise peaks with (3.3%) and without (39.5%) cis-eQTL support.

**Supplementary Figure 4.**
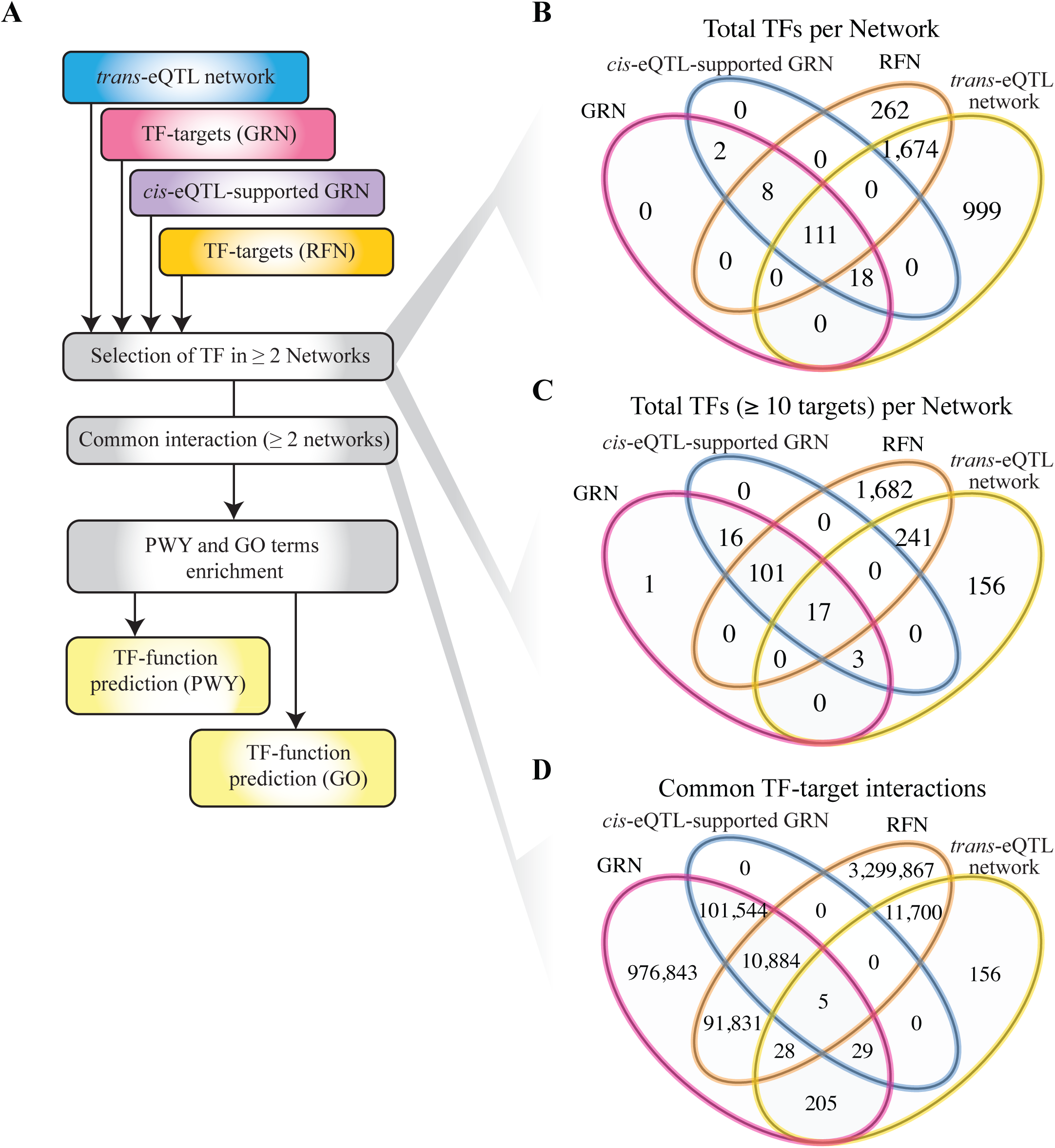
Annotating TFs by common interactions. (A) Schema of the pipeline used to annotate TFs based on target genes shared among layers (*trans*-eQTL network, GRN, *cis*-eQTL-supported GRN, and RFN). (B, C) Venn diagrams of the number of TFs shared among layers with at least one (B) and ten (C) target genes. (D) Venn diagram of the common TF–target interactions among layers.

**Supplementary Figure 5.**
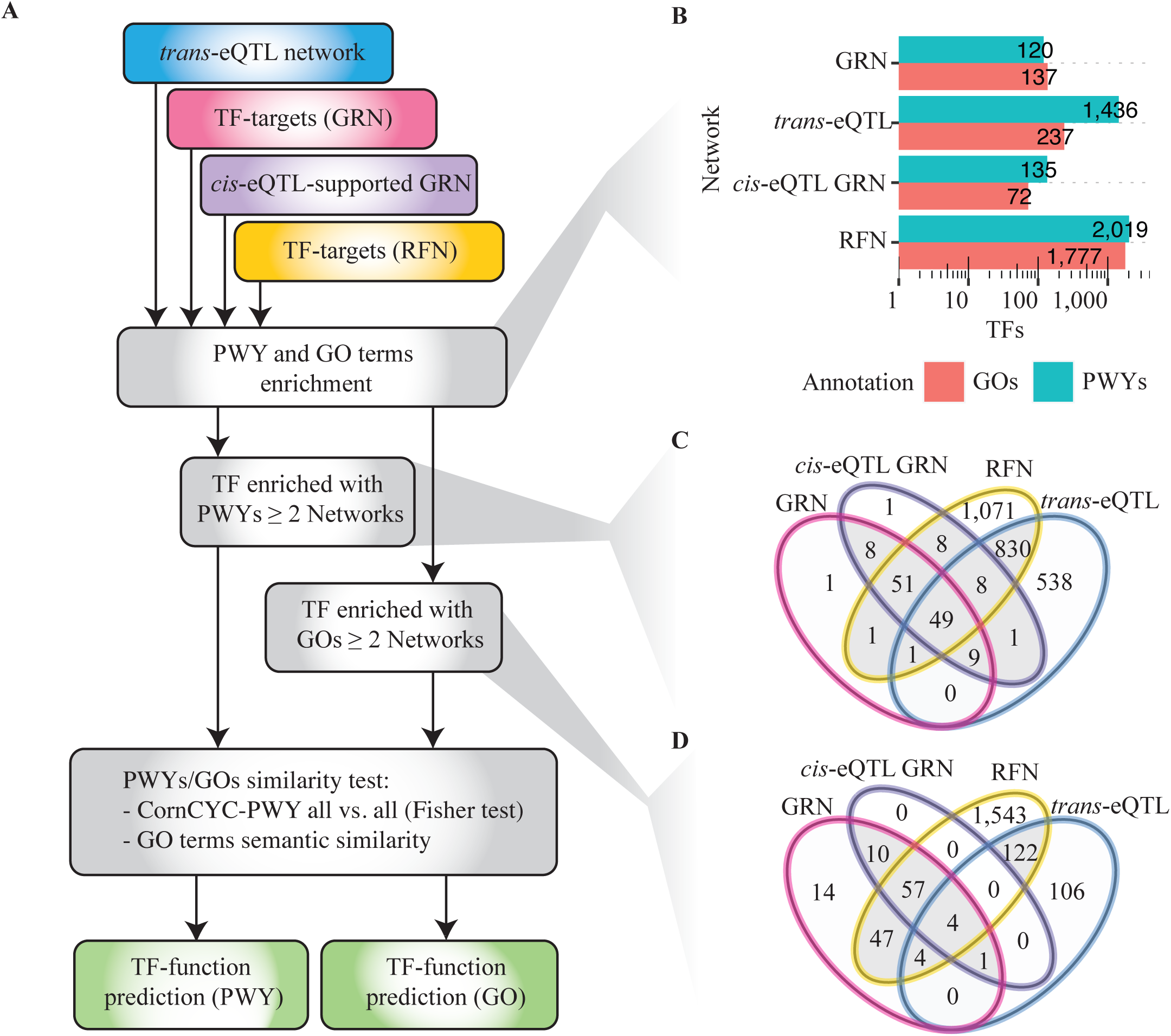
Annotating TFs by common function. (A) Schema of the pipeline used to annotate TFs based on functions shared among layers (*trans*-eQTL network, GRN, *cis*-eQTL-supported GRN, and RFN). (B) Total TFs annotated by layer and function type. (C, D) Venn diagrams of the number of TFs with at least one PWY (C) or GO term (D) commonly enriched among the corresponding layers.

**Supplementary Figure 6.**
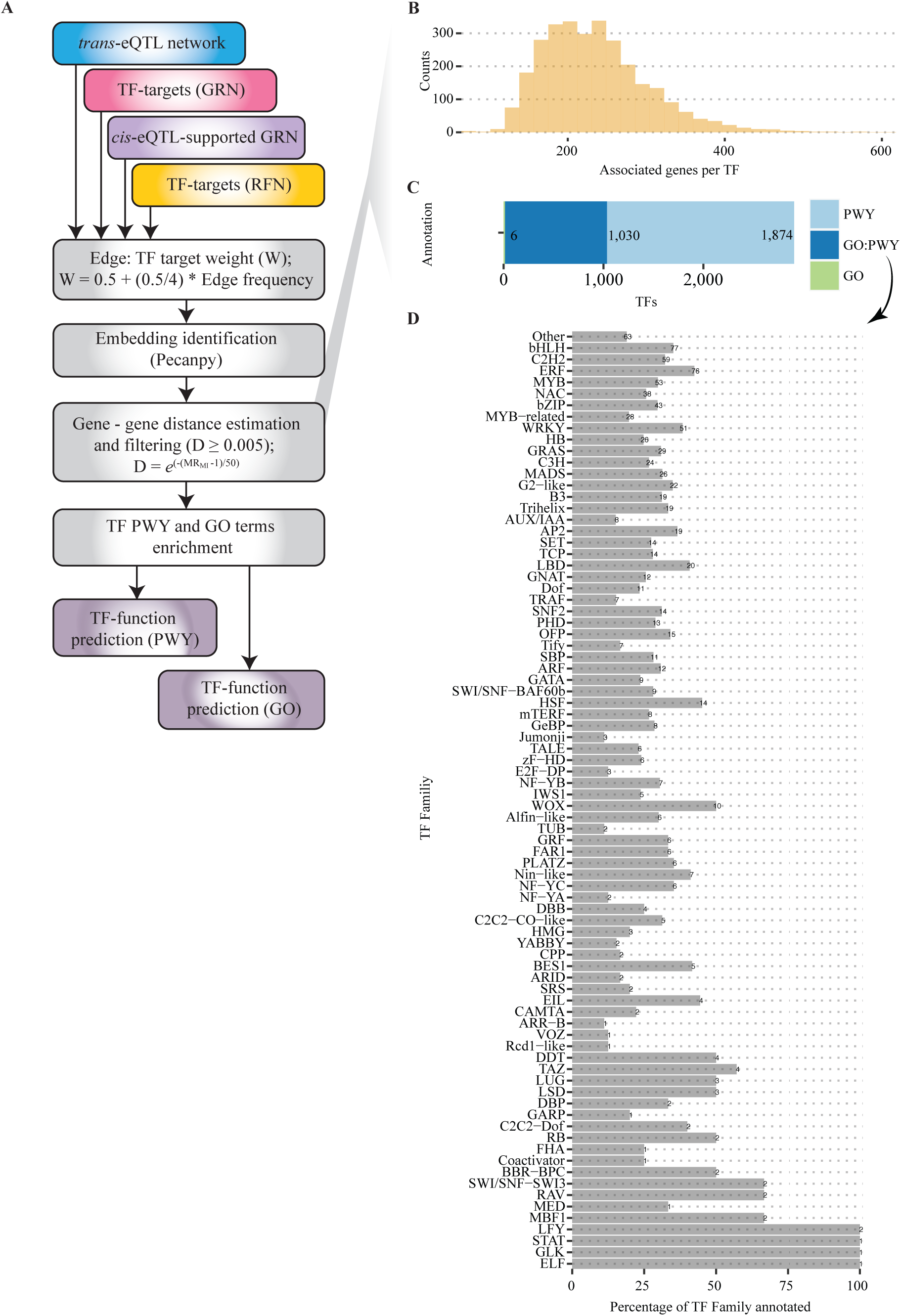
Network-based strategy for TF annotation. (A) Schema of the pipeline used to integrate the layers and identify genes with similar topological properties, defined here as network-based TF annotation. (B) Distribution of the number of genes associated per TF. (C) Total TFs annotated by enrichment with PWYs and GO terms. (D) Percentage of TFs annotated in each of the 82 TF families (and co-regulators) with at least one annotated TF.

**Supplementary Figure 7.**
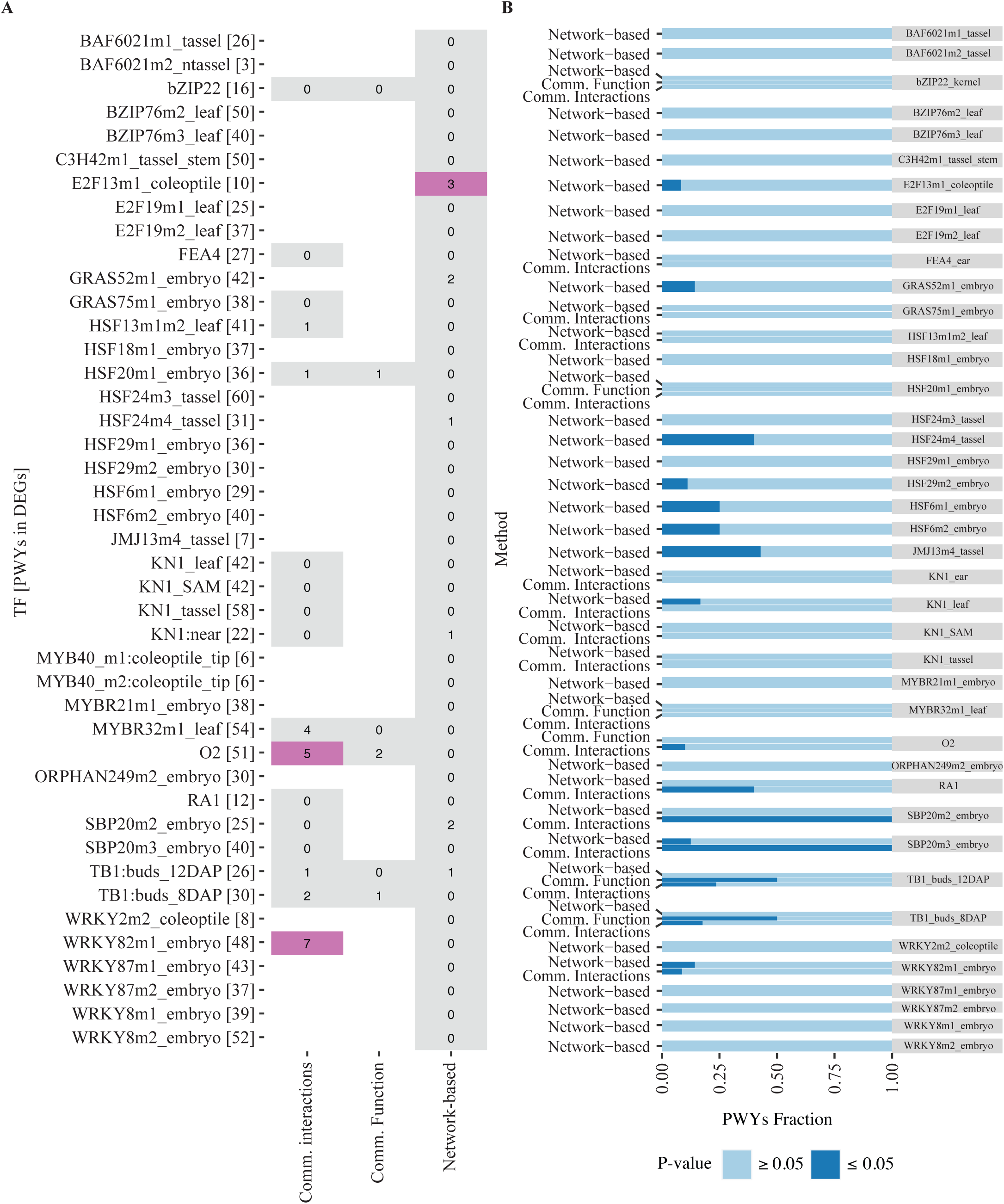
Predicted PWYs overlap poorly with knockout PWYs. (A) Heatmap of the overlap between predicted PWYs and PWYs enriched in knockouts per method. Violet boxes mark significant overlap (P ≤ 0.05, Fisher’s exact test), and white boxes denote no predicted PWYs for the corresponding TF. Square brackets indicate the number of PWYs significantly enriched in the corresponding knockout (P ≤ 0.05, Fisher’s exact test). (B) Fraction of predicted PWYs significantly enriched among differentially expressed genes per knockout assay and method.

**Supplementary Figure 8.**
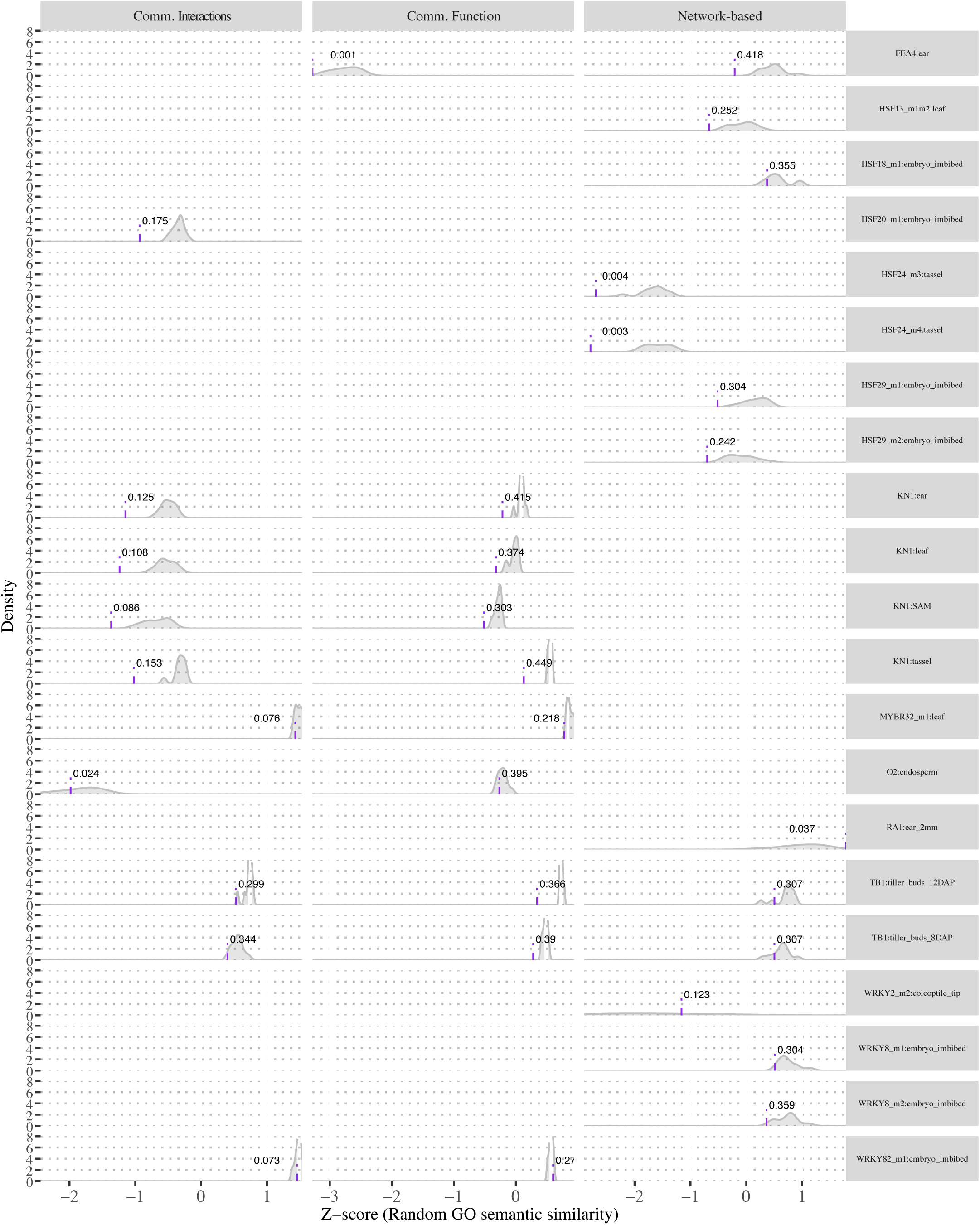
Predicted–knockout GO semantic similarity exceeds chance. Density plots of the random GO-term semantic-similarity distributions. The observed value from the real GO enrichment, with its P-value with respect to the random distribution, is indicated by the purple line.

**Supplementary Figure 9.**
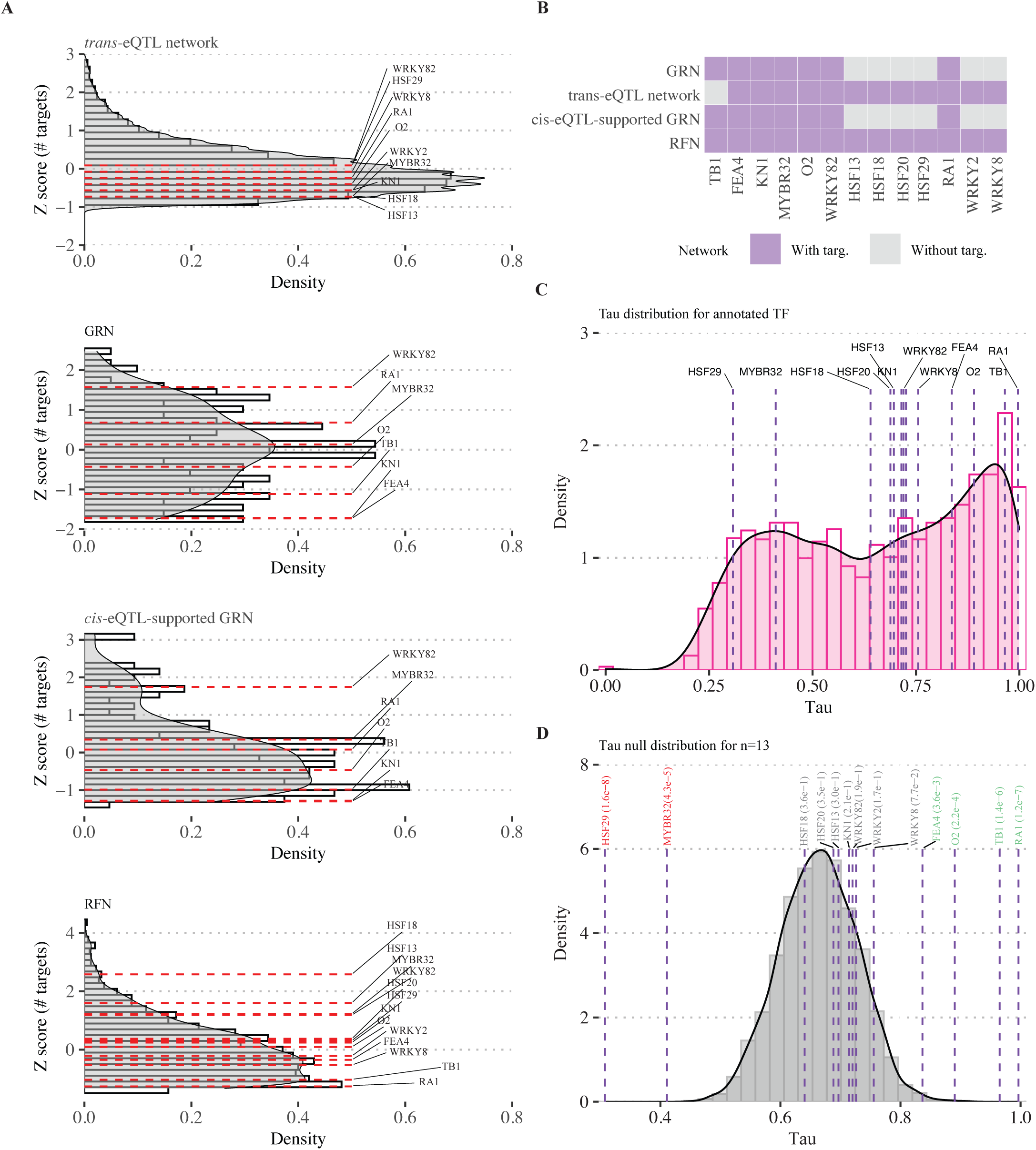
Target-gene and expression distributions of TFs vs knockout results. (A) Histogram and density plots of the scaled (Z-score) number of target genes in each layer, with the TFs used in the knockout analysis marked by dotted red lines. (B) Presence or absence of target genes in each of the four layers for every TF analyzed in the knockout assays. (C) Distribution of the Tau tissue-specificity index for the 2,910 TFs annotated with at least one PWY or GO term, with the knockout TFs marked by dotted purple lines. (D) Null distribution of Tau obtained by randomly sampling 13 TFs 1,000 times, with the knockout TFs marked. P-values were calculated against the null distribution.

**Supplementary Figure 10.**
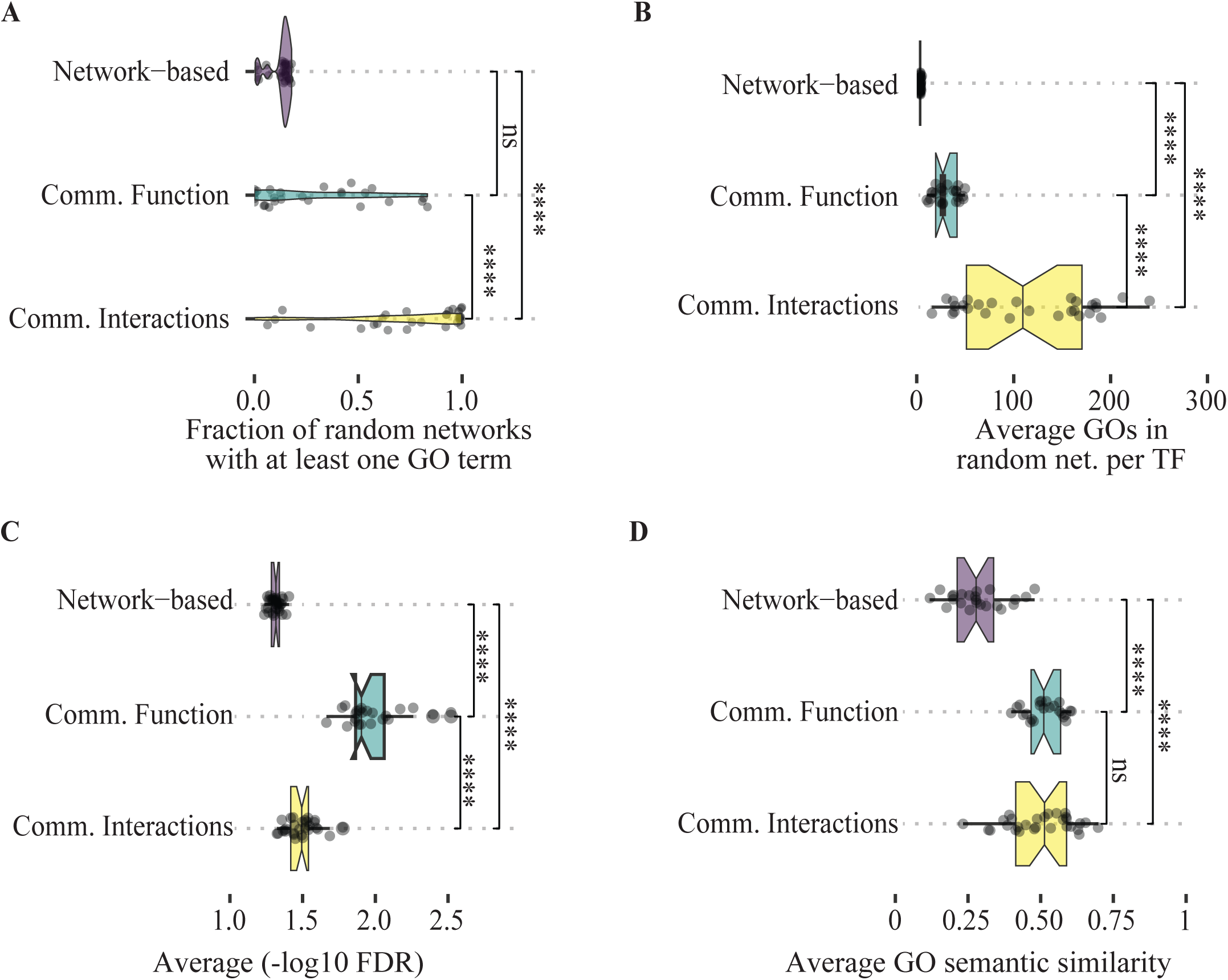
Evaluation of TF annotation against random networks. (A–D) Fraction of random networks with significantly enriched GO terms (A), average number of GO terms (B), −log10 FDR (C), and GO semantic similarity (D) observed in 3,000 random networks per method. Semantic similarity values compare each random network with the GO terms observed from the true TF–target interactions. Asterisks: *P ≤ 0.05, **P ≤ 0.01, ***P ≤ 0.001, ****P ≤ 0.0001 (two-sided t-test).

**Supplementary Figure 11.**
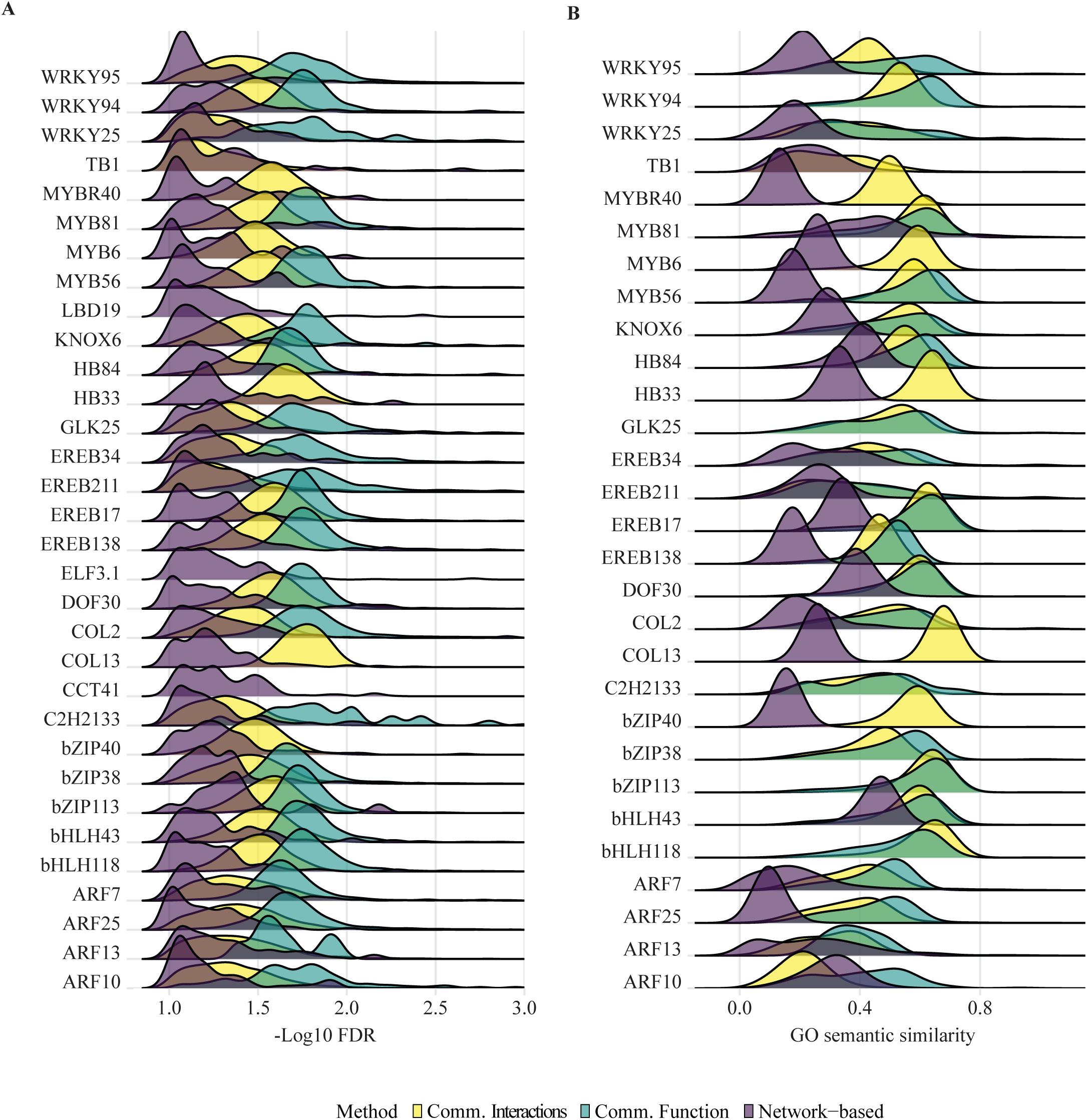
GO significance and similarity distributions from random networks per TF. (A, B) Density-ridge plots of the average −log10 FDR (A) and GO semantic similarity (B) distributions across 3,000 random networks for each TF. Semantic similarity values compare each random network with the GO terms observed from the true TF–target interactions.

**Supplementary Figure 12.**
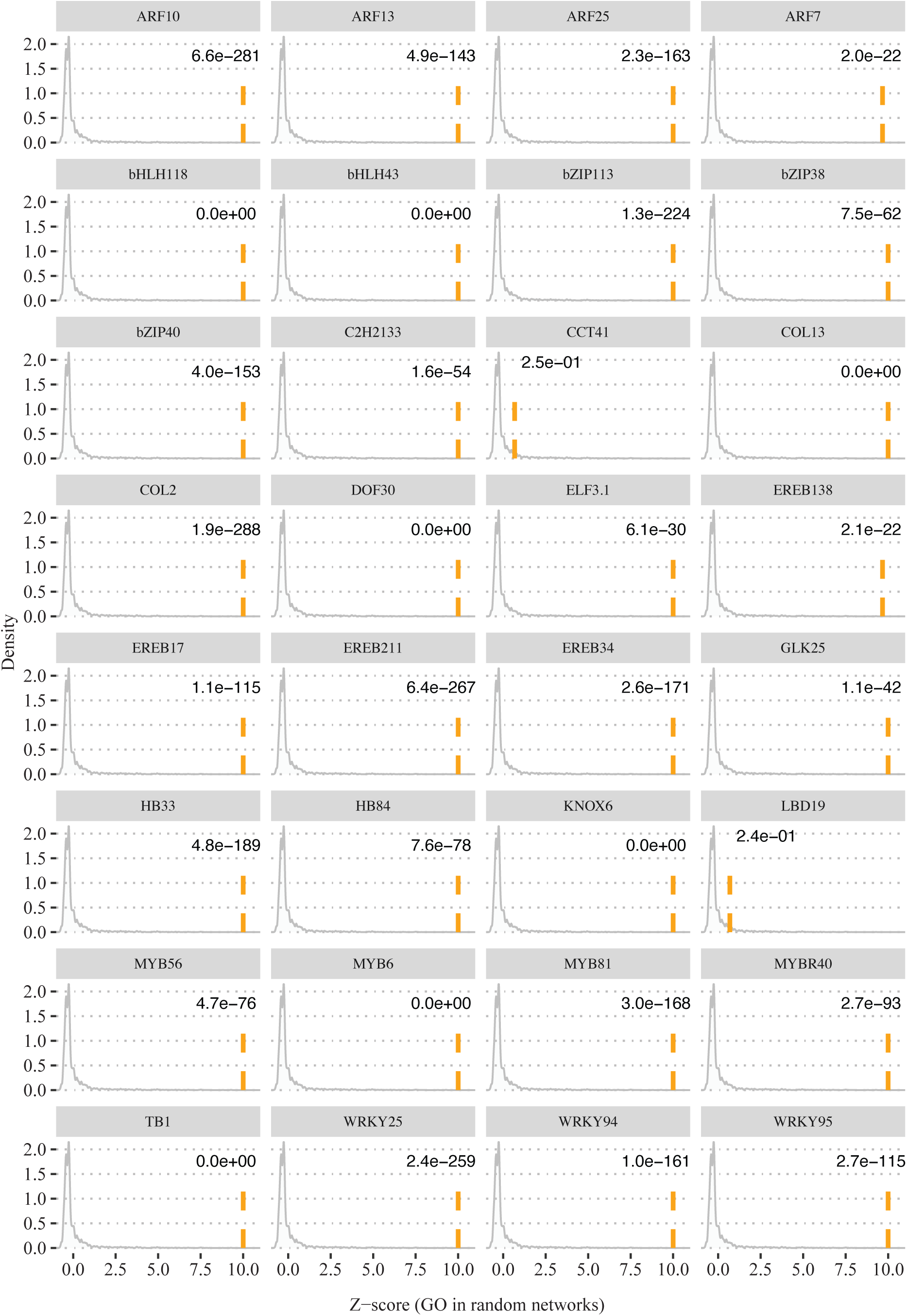
Scaled GO-term counts in random networks. Density plots of the number of GO terms predicted by the network-based method in random networks for each TF. The observed number of significantly enriched GO terms for the corresponding TF is indicated by a dashed orange line. P-values were calculated using the random distribution as the null.

**Supplementary Figure 13.**
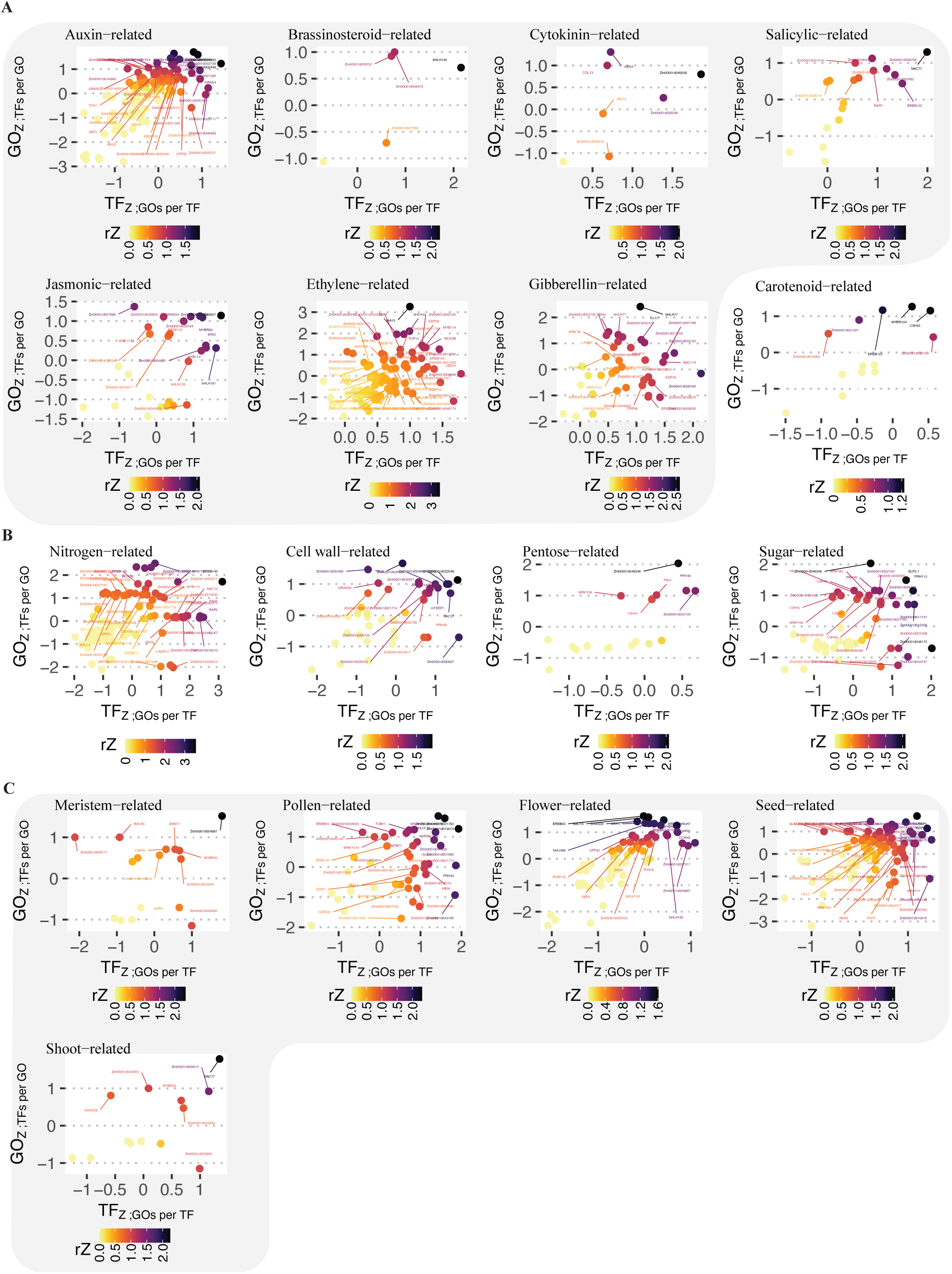
Enrichment scores for TFs and GO terms across biological processes. (A– C) Scatter plots of the reciprocal Z-scores (rZ) for hormone- (A), metabolism- (B), and development-related (C) processes.

**Supplementary Figure 14.**
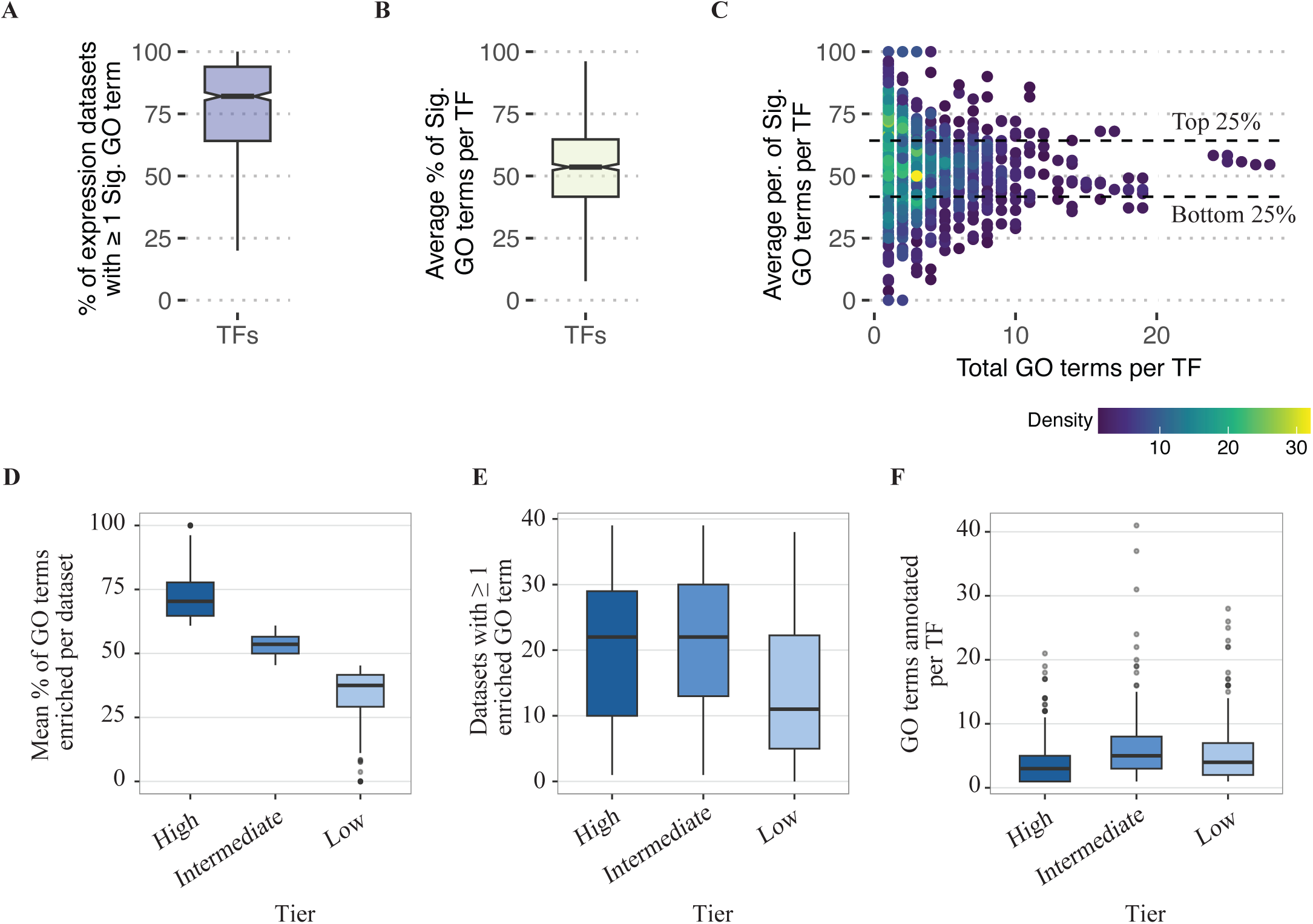
Differences in GSEA enrichment per TF tier. (A, B) Percentage of expression datasets with ≥1 enriched GO term (A) and mean percentage of annotated GO terms significantly enriched (FDR ≤ 0.1) per TF (B). (C) Total GO terms annotated per TF versus the percentage of enriched GO terms per TF, colored by TF density. (D) Average percentage of significantly enriched GO terms per tier. (E) Number of datasets with at least one significant GO term per TF and tier. (F) Number of GO terms annotated per TF, grouped by tier. Panels D–F use the tercile tiers defined in Fig 4A.

**Supplementary Figure 15.**
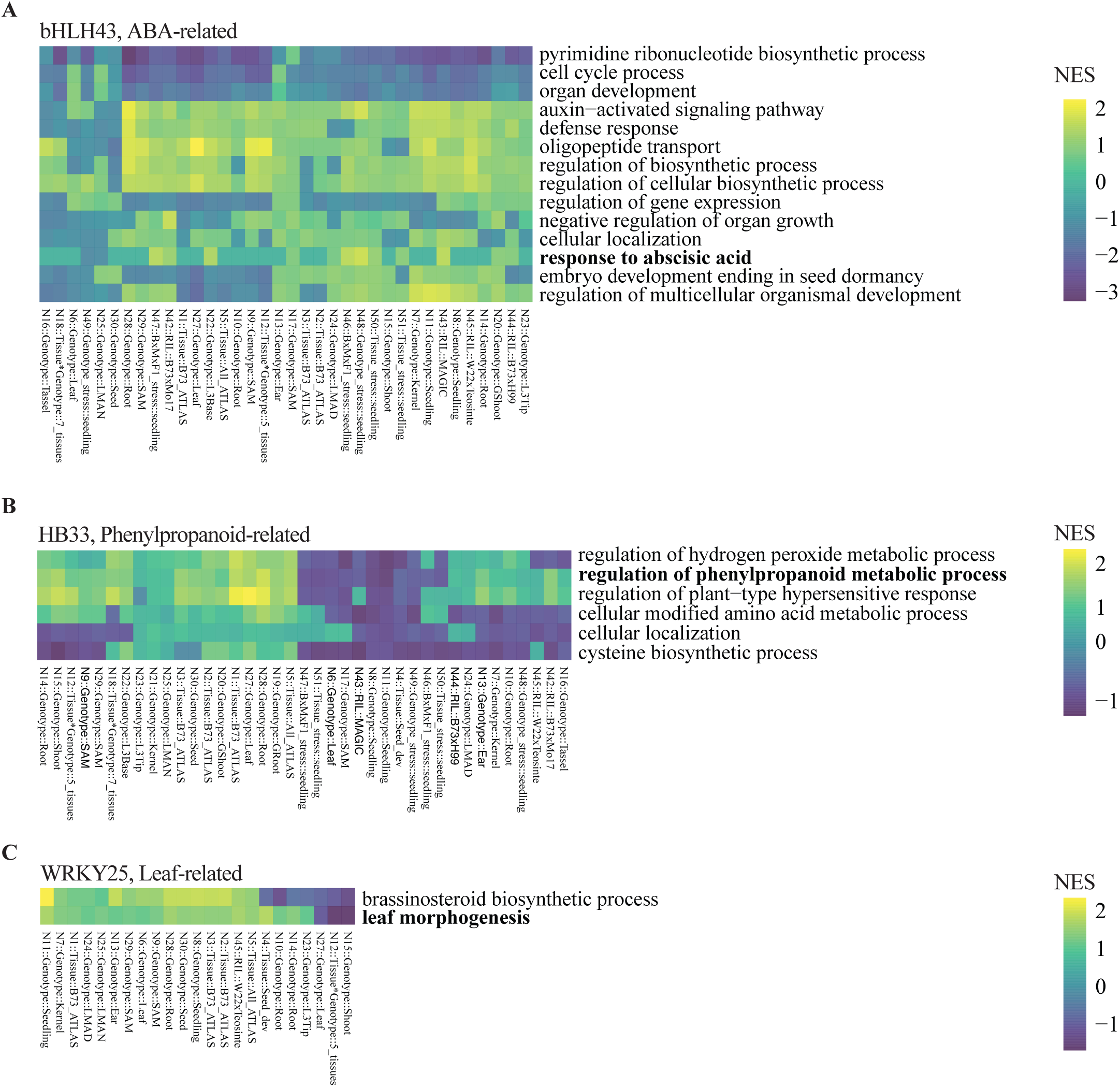
Condition-specific TF–GO regulatory effects from GSEA. (A–C) Heatmaps of the normalized enrichment score between bHLH43 (A), HB33 (B), and WRKY25 (C) and their corresponding GO terms across expression datasets.

**Supplementary Figure 16.**
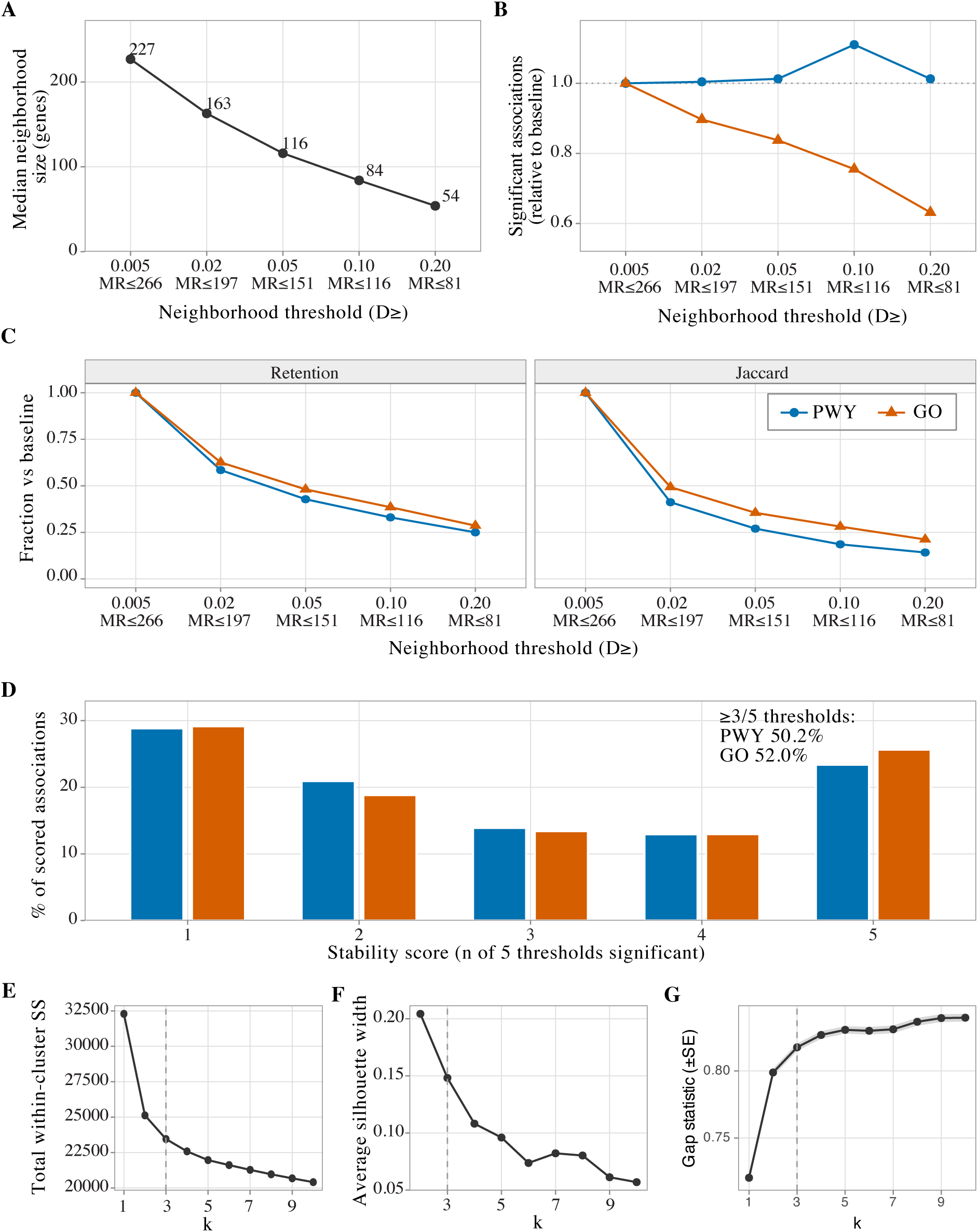
Sensitivity of network-based annotation to the embedding-similarity threshold. (A) Median functional-neighborhood size per TF at each neighborhood similarity cutoff (D ≥ 0.005 to 0.20, corresponding to mutual rank ≤ 266 to 81). (B) Number of significant TF–PWY (blue) and TF–GO (orange) associations at each threshold, relative to the baseline (D ≥ 0.005). (C) Retention of baseline associations (left) and Jaccard similarity of the significant sets (right) at each threshold, relative to the baseline. (D) Distribution of the per-association stability score (number of thresholds, out of five, at which the association is significant) for PWY and GO associations. The inset gives the fraction of associations significant at three or more thresholds. (E–G) Diagnostics for the original k-means clustering of the TF condition-enrichment profiles behind the Fig 4A tiering, total within-cluster sum of squares (E), average silhouette width (F), and gap statistic with standard error (G) for k = 1 to 10. The dashed line marks the previously used k = 3, which these diagnostics did not single out, motivating the deterministic terciles described in Supplementary File 3.

