## Supplemental files 1-3 for "Prioritizing Maize Transcriptional Regulators through Multi-Omic Network Integration"

**Supplementary Files 1–3 — Prioritizing Maize Transcriptional Regulators through Multi-Omic Network Integration**

Supplementary File 1. TF functional annotation by common interactions and common function

*Common interactions.* To identify common interactions, we compared all layers with each other (**Supplementary Figure 4A**). Summing the four layers gave ~4.6M TF–target interactions (**Supplementary Figure 4D**). As expected, the GRN and cis-eQTL-supported GRN shared the largest number of interactions (~112.5K), followed by the GRN and RFN layers (~102.7K) (**Supplementary Figure 4D**). After identifying interactions present in at least two networks, we kept 206.2K of the 4.6M interactions, comprising 934 TFs and ~20.6K target genes (**Supplementary Figure 4D**). Using target genes as a proxy for TF function, we tested the enrichment of the common target genes of each TF with PWYs and GO terms (*Methods*). Co-regulators were included in the analysis and treated without distinction from TFs. We found 11,362 significant associations between TFs and biological processes (TF–process), including 2,812 TF–PWY and 8,550 TF–GO associations (FDR $\leq$ 0.1, Fisher's exact test) (**Fig 2A**). On average, we identified ~8 PWYs and ~70 GO terms per TF (8,550 associations over 122 TFs) (**Fig 2B**). Combining the PWY and GO term results, we annotated 347 TFs, of which 225 showed enrichment only with PWYs (**Supplementary Table 8**). The remaining 122 TFs showed enrichment with both PWYs and GO terms (**Fig 2C**).

*Common function.* To identify common functions, we first tested the enrichment of target genes with PWYs and GO terms for each TF in each layer. We retained TFs with at least one PWY or GO term enriched in at least two different layers, which allowed us to explore common predictions between layers for those TFs (**Supplementary Figure 5A**). The number of enriched TFs varied across layers, ranging from 120 to 2,019 TFs for PWYs and from 72 to 1,777 TFs for GO terms. Among the layers, the cis-eQTL-supported GRN had the fewest annotations and the RFN layer the most (**Supplementary Figure 5B**). After selecting TFs with at least one PWY and/or GO enrichment, we restricted the analysis to 966 TFs (PWY) and 245 TFs (GO). For PWY annotations, the RFN and trans-eQTL network pair had the highest number of annotated TFs (888) and the GRN and trans-eQTL network pair the lowest (59). Similarly, for GO term annotations, the RFN and trans-eQTL network pair had the highest number (130 TFs) and the GRN and trans-eQTL network pair the lowest (59 TFs) (**Supplementary Figure 5C, D**). To identify common predictions at the PWY level, we evaluated gene overlap between all PWYs enriched per TF between layers ($P \leq 0.05$, Fisher's exact test) (**Supplementary Figure 5A**). A similar approach was used at the GO term level. However, given the hierarchical and redundant nature of GO terms, we used semantic similarity rather than gene overlap to determine common GO terms per TF between layers (FDR $\leq$ 0.1) (*Methods*). Together, these two annotation analyses yielded 7,081 TF–process annotations (727 TF–PWY and 6,354 TF–GO) (**Supplementary Table 8, Fig 2A**). On average, this corresponds to ~3.6 PWYs and ~58 GO terms per TF (**Fig 2B**). These associations encompass 204 TFs annotated through PWY enrichment and 110 TFs through GO term enrichment (**Fig 2C**).

Supplementary File 2. Evaluation of functional predictions against random networks

Despite recovering GO terms similar to, and enriched in, the knockout results (**Fig 2D, 2E**), we wanted to determine which method generated the fewest false positives. We therefore evaluated the identification of GO terms from ~3,000 random networks (*Methods*). We counted the number and significance of the GO terms enriched in random networks as a measure of precision, and the similarity of the observed GO terms (from true TF–target interactions) with the GO terms from random networks as a measure of accuracy. To compare predictions across methods for the same TFs, we restricted the analysis to the 32 TFs with GO predictions from all three methods (**Fig 2C**, green and blue intersection). We posited that fewer GO terms, less significant FDR values, and random-network GO terms less similar to the observed terms indicate better predictions. Consistent with our results, the network-based method identified significantly enriched GO terms in only ~12% of the random networks tested, substantially lower than the ~28% and ~72% obtained with the common function and common interactions methods, respectively (**Supplementary Figure 10A**). Concordantly, network-based predicted significantly fewer GO terms (**Supplementary Figure 10B**), with less significant P-values (**Supplementary Figures 10C, 11A**), and GO terms less similar (lowest GO semantic similarity values) to those predicted from true interactions per TF than either common function or common interactions (**Supplementary Figures 10D, 11B**), indicating that the network-based method produced the highest precision and accuracy. Notably, the common function method predicted fewer GO terms per TF than the common interactions method, with P-values of higher statistical significance and semantic similarity values on par with those of common interactions (**Supplementary Figure 10**). For any individual TF, the semantic similarity distributions of the *common function* and *common interactions* methods overlapped to a high degree, suggesting the two methods are equally noisy (**Supplementary Figure 11**).

Overall, the *network-based* method detected fewer GO terms per TF (**Fig 2B**) and had GO terms enriched in significantly fewer random networks (**Supplementary Figure 10A**). This could reflect a limited ability to identify GO term associations, as the method inherently identifies fewer GO terms per TF. To examine this possibility, we asked whether the number of GO terms observed with true interactions could be attributed to chance. For 30 of the 32 tested TFs, the total number of GO terms was significantly higher than expected by chance ($P \leq 0.05$) (**Supplementary Figure 12**). Thus, although the network-based approach yields fewer GO terms per TF, the identified terms carry biologically meaningful information that is unlikely to arise by chance alone. We conclude that, within the context of these data and layers, the network-based method is the superior approach, and we relied exclusively on network-based predictions for the subsequent analyses.

Supplementary File 3. Threshold-sensitivity analysis of network-based annotation

In the network-based strategy, each TF is annotated from its functional neighborhood, defined as the set of genes whose embedding similarity to the TF (quantified from the mutual rank of mutual information, MRMI) passes a neighborhood similarity cutoff D = e^(−(MRMI − 1)/50). The default cutoff used throughout the paper is D ≥ 0.005 (corresponding to MRMI ≤ ~266). Because neighborhood size depends on the cutoff D, we repeated the neighborhood definition and PWY enrichment analysis across five increasingly strict cutoffs, D ≥ 0.005, 0.02, 0.05, 0.10, and 0.20 (corresponding to MRMI ≤ 266, 197, 151, 116, and 81). At each threshold, the enrichment background was recomputed from the thresholded network, and the same tests and multiple-testing corrections were applied as in the main analysis.

Annotation yield was robust across this 40-fold threshold range. All 2,915 TFs retained a non-empty neighborhood at every threshold, while the median neighborhood size decreased from 227 genes at baseline to 54 at the strictest cutoff. The number of TFs with at least one significant pathway (FDR ≤ 0.05) increased modestly from 71 to 92, and the number of significant TF–PWY pairs remained between 130 and 145. The enrichment landscape changed smoothly, with the all-pairs Spearman correlation of −log10 P-values against baseline decreasing from 0.86 (D ≥ 0.02) to 0.52 (D ≥ 0.20). Turnover concentrated among marginal associations. At D ≥ 0.02, 89% of the baseline associations that lost significance still showed a positive pathway overlap, and the strongest baseline associations ($P < 10^{-5}$) were retained two to three times more often than the weakest. A robust core of 165 TF–PWY pairs across 102 TFs was significant (FDR ≤ 0.1) in at least three of the five thresholds, with 42 pairs significant in all five.

| D ≥ | MRMI ≤ | Median neighborhood | TFs annotated (FDR ≤ 0.05) | Significant TF–PWY pairs | Retention of baseline (FDR ≤ 0.1) | Jaccard vs baseline |
|---|---|---|---|---|---|---|
| 0.005 (baseline) | 266 | 227 | 71 | 131 | 1.00 | 1.00 |
| 0.02 | 197 | 163 | 79 | 130 | 0.58 | 0.41 |
| 0.05 | 151 | 116 | 89 | 145 | 0.43 | 0.27 |
| 0.10 | 116 | 84 | 91 | 142 | 0.33 | 0.19 |
| 0.20 | 81 | 54 | 92 | 144 | 0.25 | 0.14 |

To make the threshold dependence transparent in the released catalog, every TF–PWY association in Supplementary Table 8 carries a stability score, defined as the number of thresholds (out of five) at which the association is significant under the criterion used to build the catalog (raw P ≤ 0.05). Of the 23,796 catalog associations, 5,565 were significant at all five thresholds, 3,081 at four, 3,306 at three, 4,981 at two, and 6,863 at one. Half of the catalog (50.2%) is therefore supported at three or more thresholds, and we recommend prioritizing these stable associations.

Repeating the paper's exact topGO enrichment pipeline (biological process, classic Fisher) over the same five thresholds, behaved concordantly. Significant TF–GO associations (FDR ≤ 0.1 per TF) decreased from 17,596 at baseline to 11,116 at the strictest threshold, with retention of baseline associations of 0.63, 0.48, 0.39, and 0.29 across the four stricter thresholds. For the released catalog, stability was scored at the level of the Supplementary Table 8 parent terms through the recorded term-to-parent mapping. Because the Gene Ontology itself evolves, 2,253 of the 7,722 catalog associations involve terms that are obsolete or restructured in the current GO release (three years after the original analysis); these are flagged as not assessable in Supplementary Table 8 rather than scored. None of these terms reflect ontology-category mixing, as the pipeline was restricted to biological-process terms throughout. Of the 5,469 assessable associations, 52.0% were significant in at least three of five thresholds (1,403 at all five, 709 at four, and

733 at three), closely matching the stability observed for the metabolic pathway (PWY) predictions (50.2%). The stability columns in Supplementary Table 8 therefore cover all 23,796 TF–PWY associations and the 5,469 assessable TF–GO associations.

Related to the Fig 4A tiering, we also evaluated the original k-means clustering of the TF condition-enrichment profiles (829 TFs × 39 datasets). Standard diagnostics did not support k = 3, as the average silhouette width was maximal at k = 2, the gap statistic (firstSEmax, B = 100) selected k = 5, and the within-cluster sum-of-squares elbow was soft between 2 and 3. In addition, the k-means memberships were not reproducible because no random seed was recorded, and a seeded rerun on the same matrix agreed with the published memberships for only 71.5% of TFs. We therefore replaced the clusters with deterministic terciles of each TF's mean percentage of enriched GO terms (cutoffs 45.5% and 60.9%), which preserve the high/intermediate/low interpretation while being fully reproducible (Supplementary Figure 16).
